# Statistical Power or More Precise Insights into Neuro-Temporal Dynamics? Assessing the Benefits of Rapid Temporal Sampling in fMRI

**DOI:** 10.1101/2021.06.05.447164

**Authors:** Logan T. Dowdle, Geoffrey Ghose, Clark C. C. Chen, Kamil Ugurbil, Essa Yacoub, Luca Vizioli

## Abstract

Functional magnetic resonance imaging (fMRI), a non-invasive and widely used human neuroimaging method, is most known for its spatial precision. However, there is a growing interest in its temporal sensitivity. This is despite the temporal blurring of neuronal events by the blood oxygen level dependent (BOLD) signal, the peak of which lags neuronal firing by 4 to 6 seconds. Given this, the goal of this review is to answer a seemingly simple question – “What are the benefits of increased temporal sampling for fMRI?”. To answer this, we have combined fMRI data collected at multiple temporal scales, from 323 to 1000 milliseconds, with a review of both historical and contemporary temporal literature. After a brief discussion of technological developments that have rekindled interest in temporal research, we next consider the potential statistical and methodological benefits. Most importantly, we explore how fast fMRI can uncover previously unobserved neuro-temporal dynamics – effects that are entirely missed when sampling at conventional 1 to 2 second rates. With the intrinsic link between space and time in fMRI, this temporal renaissance also delivers improvements in spatial precision. Far from producing only statistical gains, the array of benefits suggest that the continued temporal work is worth the effort.

## Introduction

Functional MRI is one of the most common non-invasive brain imaging methods used to infer neuronal activity. By exploiting the coupling between neural responses and subsequent blood oxygenation changes (Logothetis et al., 2001), fMRI infers cortical activity by measuring local magnetic susceptibility changes via the Blood Oxygen Level Dependent (BOLD) signal (Ogawa et al., 1990). BOLD fMRI’s popularity is likely owed to its ease of use, high contrast to noise ratio, and the relatively high spatial precision of its functional measurements, which ranks highest amongst non-invasive in-vivo neuroim In fact, growing availability of ultra-high field magnets (e.g. >= 7T) and the development of ever more efficient acquisition software and hardware (Bollmann and Barth, 2020), it is now routinely possible to acquire functional images with submillimeter spatial resolution (e.g. 0.8 mm isotropic; (Koopmans et al., 2010; Margalit et al., 2020; Olman et al., 2012; Siero et al., 2011), or even 0.5 mm isotropic; (Vizioli et al., 2020b)), allowing the investigation of some of the most fundamental units of neural computation, such as cortical layers and columns.

Despite the second to sub-second temporal resolutions with which we can acquire images with the BOLD fMRI technique, the vascular response lags neuronal events (Logothetis et al., 2001), blurring the temporal precision relative to neuronal responses. The temporal response of the BOLD signals follows a double gamma function, with a large signal increase peaking some 5 to 6 seconds after stimulation, and a less prominent undershoot far outlasting the positive peak with an overall duration of approximately 20 seconds (Glover, 1999). These numbers differ by many orders of magnitude in comparison to the occurrence of neural events, which span the millisecond range. Consequently, fMRI’s temporal dimension has traditionally been more neglected, with the vast majority of studies relying primarily on its spatial characteristics.

This is not to say that fMRI’s temporal information has not been studied or used to interpret neural activity. For example, in considering relative temporal differences on how stimuli and task demands drive differential responses, there have been a number of investigations, including how delayed decision making affects the BOLD signal (McGuire and Kable, 2015), investigations into the variability in temporal dynamics in the non-human primate auditory pathways (Baumann et al., 2010), or into the variability in the context of visual and semantic processing (Avossa et al., 2003; Formisano and Goebel, 2003; Vu et al., 2016).

Substantial research efforts have produced more efficient hardware and software, decreasing fMRI’s repetition time (i.e. the time required to record a whole volume; TR). Recently, using these highly accelerated acquisition protocols, researchers have been able to acquire BOLD time series with unprecedented whole brain temporal resolutions (i.e. ∼300-500 ms; Lewis et al., 2016; Vu et al., 2016). However, as most of what we know about the temporal characteristics of the BOLD signal have been learned with supra-second temporal resolutions, the impact of such high temporal resolution remains to be determined. Further work is required to understand the full potential of faster sampling in enabling more precise insights into neuro-temporal dynamics, more accurate characterization of the BOLD response function, or potential gains in statistical power compared to more conventional temporal resolutions. With faster TRs and improved signal to noise ratios (SNR) there is the exciting possibility of uncovering faster vascular dynamics, such as the initial dip (Menon et al., 1995).

In this work we aim to quantify the benefits of acquiring BOLD fMRI data with sub-second temporal resolutions. While the bulk of this work will focus on task-based fMRI, we will also touch upon the benefits of temporal resolution in the context of resting state fMRI, as resting state analyses are increasingly popular and widely used. We will review the literature and complement existing findings with empirical data to demonstrate the impact fast acquisitions have with regards to spatial and temporal information in fMRI. We will briefly touch upon the technical development that rendered fast (here defined as <1s) and ultra-fast (here defined as <0.5s) fMRI possible. We will also examine the challenges associated with collecting, analyzing and interpreting fast data and consider the cost vs. benefit trade-offs of rapid data acquisition. Finally, as the BOLD signal in space and time are intrinsically linked and precision in one domain could inform or give rise to precision in the other, we will examine whether the ability to acquire higher temporal resolutions allows us to simultaneously and more precisely consider the information content from both dimensions.

### Early temporal work

Interest in fMRI began very early, shortly after its inception, and built upon the prior literature of mental chronometry, which primarily used reaction time measures to examine the timing of cognitive processes (Posner, 1978). Despite temporal lag and blurring, the goal for fMRI in this context is to determine the relative order of neuronal events within and across brain regions. While it is likely to remain unattainable to precisely identify the absolute ordering or directionality of neuronal events with fMRI, uncovering the relative timing of these events is more within reach. Historical examinations of the BOLD signal noted that events with an 8 second interstimulus interval (ISI) were visibly separate in signal traces (Bandettini et al., 1993). Another study (Kim et al., 1997) was able to resolve 2s long blocks of finger tapping tasks separated by only 3 seconds. These early proof-of-principle studies showed that the BOLD signal could be used to examine temporal dynamics, but such slow experimental designs were not optimized for widespread usage.

Following developments in statistical analyses and fMRI methods, work in mental chronometry grew more advanced. One method, used by Menon and colleagues, involves performing a linear fit to the initial rising portion of the hemodynamic response (HDR; (Menon et al., 1998)). The point at which this best fit line intercepts the baseline BOLD signal is known as the time-to-onset (TTO). By comparing this onset time between regions, they uncovered latency differences between the primary visual cortex and the supplemental motor area in a visually cued motor task (Menon et al., 1998). Though the approach put forward by Menon et al. became a popular method for determining the relative timing of events, alternative approaches, such as fitting a single gamma function and deriving its onset (TTO) or the time-to-peak (TTP) (Miezin et al., 2000) have also been used. In general, these methods take advantage of the characteristic shape and relative consistency of the HDR in order to deconvolve sub-TR variability in neural timing. Studies like these, however, often implemented very specific task designs, which are not broadly applicable or used substantially less than whole brain coverage in order to reduce the TR.

Later work attempted to provide a more comprehensive view of BOLD temporal effects, using complex sequential tasks (combining auditory perception, mental imagery, and motor responses) to measure the order of mental operations across larger areas of the cortex (Formisano et al., 2002). Formisano et al. (2002) for example were able to detect sub-TR and sub-second delays between multiple regions despite TRs on the order of 2 seconds. These studies show that even with longer TRs, it is possible to extract rich temporal information from BOLD responses due to the consistent nature of the HDR. Of course, this consistency only emerges under specific circumstances, that is, within the same area, in response to the same or very similar stimulus. Another method to derive sub-TR timing effects that builds on this consistency combines the canonical HDR with its temporal derivative in a general linear model (GLM) framework. By comparing the amplitude of the temporal derivative to the canonical response, one can determine the relative timing of events (Henson et al., 2002; Liao et al., 2002). Building on temporally shifted HDRs, an alternative approach entails evaluating model fits between the fMRI time signal and HDRs with different time lags, simulating different timings of neuronal events. This approach has been argued to potentially lead to a temporal precision spanning the 200ms (Hernandez et al., 2002) or even 100ms (Bellgowan et al., 2003) range. In an alternative approach, Sigman and colleagues manipulated the relative timing and duration of a series of sequential events to jointly consider the magnitude and phase of the BOLD response in order to determine timing with 100ms precision (Sigman et al., 2007). By taking advantage of repetition suppression effects, Ogawa and colleagues showed that fMRI could even detect modulations in the 10s of milliseconds differences when considering cross hemispheric inhibition or neural refractory processes due to short interstimulus intervals (Ogawa et al., 2000).

These studies highlight early interest in fMRI temporal dimension. However, in spite of these dramatic temporal findings, early reports present a number of limitations. These approaches are in fact limited in scope, as they are only applicable to a small subset of possible scientific contexts and questions. This limitation is related to a number of factors, including the necessarily constrained or highly specific experimental designs, the typically large voxel sizes, or very limited fields of view. As such, a clear need emerged for methods that allowed for faster sampling in conjunction with whole brain acquisition or, alternatively, a limited field of view with substantially higher spatial resolution.

### Advancements in MRI

To date, most of the fMRI applications looking to exploit temporal information did not have the benefits of fast or sub-second temporal sampling combined with relatively high SNR/CNR and large, or even whole, brain coverage (e.g. de Zwart et al., 2005; Weilke et al., 2001). In recent years there have been substantial improvements in magnet hardware, including an increasing prevalence of ultrahigh field (>= 7 Tesla) strengths and improvements in transmit (Wu et al., 2019) and receive coil arrays (Uğurbil et al., 2019). The use of higher magnetic fields results in improved image SNR as well as increases in the CNR of the BOLD fMRI signal (De Martino et al., 2018; T. Vu et al., 2017). These gains can be traded in for increased spatial resolution (as has historically been the case), increased temporal resolution, or some combination of both. The development in coil array technology has permitted the spatial encoding of signals via the sensitivity profile of the individual coils, with higher channel count coils allowing for more precise spatial encoding (Uğurbil et al., 2019). Spatial encoding using information from the RF coil arrays means less required spatial encoding from MRI gradient coils or ultimately the feasibility of accelerating conventional image acquisitions which rely on gradient encoding alone. Multi-channel array coils can be used to spatially encode information during the encoding of a single slice, which primarily facilitates the acquisition of higher spatial resolutions, or used to encode multiple slices simultaneously, which can significantly improve the temporal resolution of the acquisition (Larkman et al., 2001; Moeller et al., 2010).

Simultaneous multi-slice (SMS) or multiband (MB) imaging (Breuer et al., 2005; Feinberg et al., 2010; Larkman et al., 2001; Moeller et al., 2010) was introduced to reduce the total time needed to acquire a volume by collecting multiple slices simultaneously. This led to a revolution in the fMRI field for multiple reasons. While introduced to improve the temporal resolution or efficiency, this did not mean that it was only relevant to high temporal resolution applications. On the contrary, since higher spatial resolution studies were inherently temporally inefficient due to the many more slices required, the introduction of MB/SMS actually made higher spatial resolution studies feasible via much more tolerable TRs or allowing for increased coverage in field of view limited studies. The method is, of course, equally amenable to improving the temporal sampling for a given spatial resolution and coverage. By maintaining a constant resolution and coverage, entire volumes can be acquired in a fraction of the time – up to 6 or even 10 times faster, as was demonstrated by the Human Connectome Project (Uğurbil et al., 2013). Alternatively, MB has been exploited by auditory researchers to allow more “dead time” between TRs, enabling longer stimulus presentation time in the absence of hardware related acoustical noise (De Martino et al., 2015). With typical 3T fMRI voxel resolutions (i.e. 2-3mm isotropic), it is possible to get whole brain images in less than a second or even under 500ms. Novel acceleration methods continue to be developed, pushing acquisition times for reasonable brain volumes well below 500 or 100 ms (Chang et al., 2013; Lin et al., 2011; Vu et al., 2018). Collectively, the increase in receiver coil elements, improved accelerated acquisition and reconstruction methods (Koopmans and Pfaffenrot, 2021), and higher field strengths, all provide an encouraging outlook for rapid temporal sampling in fMRI. However, as temporal resolution becomes finer, appropriate analytical strategies to deal with these super-fast acquisitions are required.

## Statistical and Methodological Considerations

### Hemodynamic response variability

The conventional HDR has served the fMRI community well and, through parametric manipulations and analyses strategies, it is possible to determine sub-TR shifts, even with relatively low temporal resolution. However, the shortcomings of the canonical HDR, derived from sensorimotor and auditory cortices at 1.5 Tesla with 2.2×2.2×5 mm^3^ voxels (Glover, 1999), are becoming increasingly clear. It has long been known that the HDR can vary depending on the duration of the stimulus (Boynton et al., 1996; Glover, 1999), the precise stimulus presented (Boynton et al., 1996; Thompson et al., 2014), and the specific region studied (Gonzalez-Castillo et al., 2012; Handwerker et al., 2004; Taylor et al., 2018). Statistical approaches, such as adding more basis functions to the conventional HRF to account for temporal shifts and dispersion (Friston et al., 1998; Henson et al., 2001; Pernet, 2014) have improved, but not eliminated, the impact of hemodynamic variability at typical sampling rates. Rapid sampling can also mitigate these effects, allowing the HDR to be better characterized (Lewis et al., 2018; Lin et al., 2018, 2013). Unbiased estimates of the HDR can also uncover elusive temporal features, such as the initial dip (Buxton, 2001), which is absent from canonical HDR models.

In modeling the HDR with insufficiently flexible basis functions, or inadequate temporal resolution, temporal differences can instead manifest as differences in amplitude, which can lead to incorrect inferences (Lindquist et al., 2009). This challenge increases as briefer stimuli (i.e. less than 1 second) are used. When stimuli are 1 second or longer, the BOLD response tends to add linearly, such that a response to a 2 second stimulus matches the summation of a pair of 1 second stimuli (Boynton et al., 2012, 1996). With rapid sampling and brief stimuli, the hemodynamic model requires further consideration and models may need to be updated to account for these differences. A discussion of the precise shape of the HDR under all possible stimulus types, durations and sampling regimes is beyond the scope of this paper (see Polimeni & Lewis in the current issue for further consideration of these points: Polimeni, and Lewis, 2021).

A potential pitfall for any indirect measure of neural response such as fMRI is the difficulty in differentiating between vascular and neuronal temporal dynamics (Bandettini, 2014; Drew, 2019). For example, there is the danger of interpreting vascular delays as meaningful temporal differences in neuronal responses. Findings that longer time-to-peak is correlated with larger FWHM responses appear to uncover venous effects that do not necessarily hold neuronal basis (de Zwart et al., 2005). That is, hemodynamic signaling delays and blood transit time lead to larger and sluggish responses in the superficial venous drainage, a factor that should be considered when averaging within regions of interest for example. As such, extracting neuronally relevant temporal information from BOLD fMRI data can be challenging.

Regional variability, as measured by breath holding induced hypercapnia experiments, suggest that vascular delays vary across the human brain (i.e. up to +-5 seconds; Chang et al., 2008; Moia et al., 2021), though this range values can reflect draining veins, such as the superior sagittal sinus (Chang et al., 2008). These effects can be measured, and potentially accounted for to reduce their deleterious impact (Erdoğan et al., 2016). Moreover, this variability is highly regional and experiments with reversed stimulus orderings suggest that fMRI can successfully capture interregional latency differences (Lin et al., 2013). Alternatively, one can examine ipsi- and contralateral activation, as the approximate left/right symmetry of the brain yields similar responses at zero delay (Yeh et al., 2013) (though there is of course variability at finer scale Lin et al., 2018; Park et al., 2019).

Beyond these effects, care must be taken to distinguish between delays due to the processing of stimuli within a region and delays in processing between regions. For example, even in a relatively simple visually cued button response task, there are a number of latencies to account for: the latency between visual input and neural firing in the visual cortex (∼30ms), the onset of detectable BOLD signal changes (∼2000ms), the conduction and processing time between V1 and the SMA and on to motor cortex, as well as the conduction time from M1 to the appropriate muscle groups (∼120ms) (Menon et al., 1998). Additional latency differences can occur as stimuli take longer to process and identify, revealing putative increases in neural processing (Bellgowan et al., 2003).

While optimal experimental design for such techniques is beyond the scope of this manuscript, estimation of the HDR, and thus its actual parameters such as onset time and time-to-peak, is improved using short stimuli with jittered interstimulus intervals (Birn et al., 2002; Dale, 1999). In addition to improving estimations of the underlying HDR, there appear to be other reasons to prefer brief stimuli (<=1s), as these may be associated with fewer vascular artifacts and greater temporal precision (Hu et al., 1997; Menon and Kim, 1999). Analyses appear to benefit from a focus on a consistent onset and rise (such as time to half) of the fMRI response, as other measures such as time to peak or the post-peak portion of the response may reflect contributions from ongoing or prolonged activity, rather than initial response to the stimulus (Bellgowan et al., 2003; Menon et al., 1998). Alternatively, differentiating the timing of neuronal processing across tasks and or stimuli can be performed in the typical GLM framework by using a canonical response in addition to derivatives (Henson et al., 2002).

### Statistical implications of faster TRs in fMRI

Shorter TRs can extract timing information of cognitive processes (such as face processing (Gentile et al., 2017) and can lead to more precise parameter estimates, higher tSNR, and larger t-statistics (G. Chen et al., 2015; McDowell and Carmichael, 2019; Posse et al., 2012), which can be loosely defined as the estimated amplitude of the BOLD response (or contrast) divided by the model’s standard error. Other works find more modest effects, with limited benefits due to faster sampling (Bhandari et al., 2020; Demetriou et al., 2018). These mixed results may suggest that faster sampling combined with typical analysis methods are unlikely to give large benefits, however this should not be a cause for concern. The primary issue at hand is that most comparisons between conventional BOLD fMRI and faster sampling MB based BOLD fMRI have not considered an exhaustive comparison of how the removal of structured noise (discussed below), could improve results. In addition, these papers use the canonical HDR rather than customizing the response for each participant, task or voxel. Finally, these papers consider, by and large, typical analyses strategies examining groups of subjects. While these analysis strategies are valid and widely used, they may miss or average out the effects that are most interest to rapid fMRI researchers. Unsurprisingly, the benefits of faster sampling may require additional processing steps or special considerations to fully gain the benefits or appropriately handle the corresponding tradeoffs.

The increase in temporal samples is the most obvious benefit of faster sampling, which is typically equated with a one-to-one increase in degrees of freedom. However, individual volumes of fMRI data are not statistically independent, and instead exhibit a strong temporal autocorrelation, which must be accounted for in order to produce valid inferences (Bullmore et al., 1996) when using the typical t-statistic/p-value approach. Failure to do so will result in an artificial inflation of statistical power, especially for GLM related contrasts and associated p-values. In conventional fMRI, this problem is often addressed using relatively simple autoregressive models (i.e. AR(1)) (Friston et al., 2002), which attempt to estimate the amount of autocorrelation in the timeseries using only one preceding timepoint. While this appears to be valid for 2 to 3 second sampling rates, it is inadequate for second to sub-second TRs, leading to inflated t-statistics (see Figure 1D) and concerns about false positives (Bollmann et al., 2018; Chen et al., 2019; Lenoski et al., 2008; Olszowy et al., 2019; Purdon and Weisskoff, 1998). On the other hand, beta values, reflecting BOLD response amplitude, are minimally changed by accounting for temporal autocorrelations (see Figure 1C). As sampling rates increase, the relative contribution of thermal noise increases (due to lower SNR from shorter TRs) and autocorrelation effects span more volumes (Chen et al., 2019). Modeling physiological noise using methods such as RETROICOR (Glover et al., 2000; Olszowy et al., 2019), can lead to whiter residuals, however it does not fully solve the autocorrelation issue (Bollmann et al., 2018), so these more advanced methods are still required.

**Figure 1.**
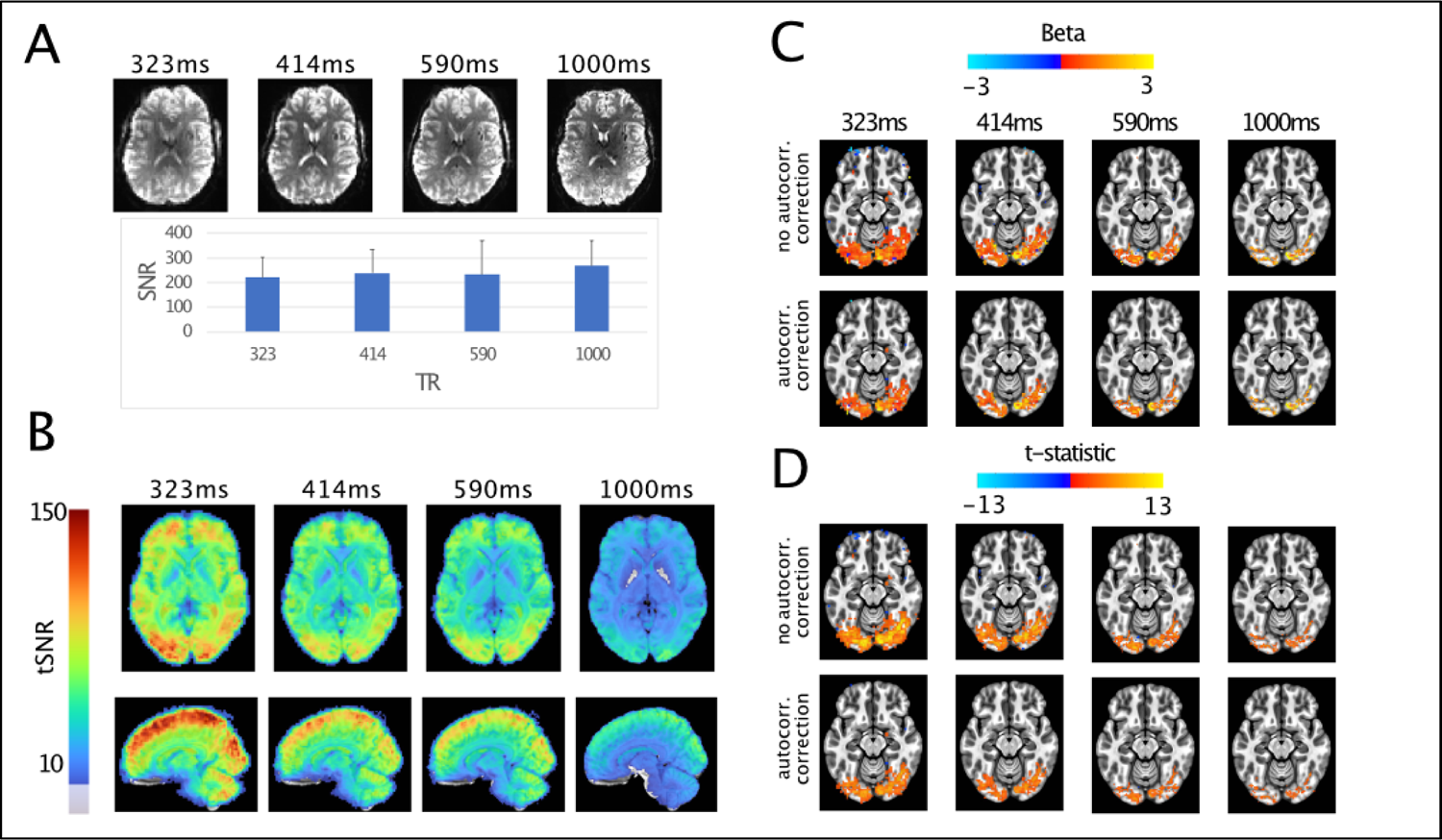
Effects of faster sampling on statistical and signal characteristics. A) Image SNR characteristics, with images from a single subject. Image SNR values were comparable across sequences (with largely overlapping error bars representing the standard errors across subjects), with a slight bias towards the 1000 ms protocol. B) tSNR measures, average over subjects. Faster sampling leads to substantial gains in tSNR throughout the brain, with the largest effects visible with a sampling rate of 323ms. C) Activation amplitudes (betas) for the nonzero phase face stimuli compared to baseline. Betas are little changed due to accounting for autocorrelation in the timeseries. D) T-statistics for the same contrast. Unlike betas, both the extent and the magnitude of the t-statistics is reduced when we attempt to account for temporal autocorrelation. These effects are most pronounced for the 323ms data. For visualization purposes, the beta and t-statistic maps use a t threshold of p<0.01 from the average t-statistics.

In dealing with autocorrelation correction, estimating parameters on a voxel by voxel or tissue by tissue basis is preferred for accuracy (Eklund, Anders et al., 2012; Lenoski et al., 2008) This is due to variability in the amount of temporal autocorrelation across the brain (Kaneoke et al., 2012), particularly between grey and white matter (Worsley et al., 2002). While the basic AR(1) is insufficient, there is not a clear answer as to what model order is required. AFNI uses a voxel-wise autoregressive moving average (ARMA(1,1)) (Chen et al., 2012), which combines the AR(1) model with a moving average (MA(1)) component to account for the presence of thermal noise in the fMRI timeseries. This relatively simple addition appears to perform well, even at sampling rates as low as 645 milliseconds (Olszowy et al., 2019), though there is potential for improvement.

In contrast to the default AR(1) model, SPM has recently implemented a method called FAST, which uses a collection of exponentially decaying functions and their derivatives to fit the timeseries autocorrelation, which also successfully reduces false positives (Corbin et al., 2018; Olszowy et al., 2019). Fitting complex AR models for each voxel does raise statistical as well as computational concerns. While higher AR models (i.e. considering more time lags) can be used to effectively account for autocorrelation, estimating higher and higher model orders is computationally inefficient and may result in overfitting, leading to spurious estimates (Bollmann et al., 2018). A recently developed approach nicely handles both the voxel-by-voxel fitting and excessive model order concerns by using methods based in information theory to select the autoregressive model order, rather than using a modification of the AR(1) approach (Luo et al., 2020). Tested against resting state data with 300 and 500ms TRs as well as simulated data they show that this method successfully controls false positives, though model orders as high as AR(10) may be required.

Collectively these findings show that, as sampling rates increase, the field should consider autocorrelation approaches that estimate use higher model orders and perform estimates on a voxel-by-voxel basis. These concerns, which primarily relate to conventional GLM-based approaches for statistical inference, apply not only to fast TR data, but to fMRI in general, and become more apparent in extreme cases, such as that of ultra-fast fMRI.\

One frequent approach, equally applicable to standard and fast fMRI, that effectively circumvents the above considerations related to the inflation of statistical power, is the implementation of independent 2^nd^ order inferential statistical tests. When carried out across sufficiently large samples, straightforward group inferences may be minimally affected by inflated single subject statistics (Bhandari et al., 2020; Kirilina et al., 2016). For other cases, more advanced analyses can be performed at the within-subjects level, across, for example, single runs estimates of BOLD responses, which may benefit from denser time course sampling.

Univariate (e.g. Dowdle et al., 2021) and multivariate (e.g. Kriegeskorte and Bandettini, 2007; Vizioli et al., 14 2020a) parametric tests using subjects or runs as independent observations are in fact routinely used.

Nonparametric statistical approaches, such as bootstrap confidence interval that make fewer assumptions about the data distribution, thus avoiding many of these concerns, are also a valid alternative. In other words, the concerns about inflation of statistical power with ultra-fast fMRI are primarily rooted in GLM-based statistical inferences. These concerns can be mitigated by assigning statistical significance on the basis of empirical p-values that are derived from independent estimates of BOLD activation, such as different subjects or experimental runs, rather than using GLM contrast-based p-values. When considering statistics derived from separate runs for example, it is also more straightforward to examine individual differences in space and time, which are necessarily blurred when many subjects are combined.

In summary, fMRI’s increased temporal resolution, which allows a better characterization of BOLD responses, also leads to a significant increase in the number of temporal samples, introducing an additional degree of complexity in analyzing and interpreting ultra-fast data. However, there are a number of more or less refined strategies to deal with these additional complexities, that can also be circumvented using non GLM-based inferential statistic, allowing exploitation of fast-TR data.

### Empirical effects of faster sampling rates

Though a number of questions remain about the best methods to use and how acquisition strategies will alter metrics such as tSNR, some of these effects can be readily demonstrated. Here we show evidence of these statistical considerations with a dataset collected at 7T, using visual stimuli consisting of faces at varying phase coherence levels.

Four (2 female) healthy right-handed subjects (age range: 18-31) participated in the study. All subjects had normal, or corrected vision and provided written informed consent. The local IRB at the University of Minnesota approved the experiments. In the scanner, participants were instructed to perform 2 tasks: face detection and fixation. In the former, we varied the phase coherence of the face stimulus from 0% to 40% coherence (10% step size) and participants were instructed to respond as quickly as possible by pressing one of 2 buttons on a button box to indicate whether they perceived a face. In the latter, participants were instructed to press one of the 2 buttons on the button box when the fixation color changed to red. Every 500 ms, the fixation changed to one of five colors (red, blue, green, yellow or cyan) in a pseudorandom fashion, avoiding consecutive presentations of the same color. The frequency of button presses was kept constant across tasks. Visual stimuli were identical across tasks in order to examine differences based only on attentional effects. Tasks were blocked by run and counterbalanced across participants.

We acquired 3 runs per task. Each run lasted approximately 3 mins and 22 secs and began and ended with a 12-second fixation period. Within each run, we showed 40 images (5 phase coherence levels x 4 identities x 2 genders) presented for 2000 ms, with a 2000 ms interstimulus interval (ISI). Importantly, we introduced 10% blank trials (i.e., 4000 ms of fixation period) randomly interspersed amongst the 40 images, effectively jittering the ISI.

All functional MRI data were collected with a 7T Siemens Magnetom System using a 1 by 32-channel NOVA head coil. BOLD fMRI data were collected using 4 unique sequences varying in TR from 1000 to 323ms (Table 1). For each sequence, parameters were adjusted, including spatial resolution, in order to produce images with comparable signal to noise ratios (SNR) despite changes in TR. The flip angle was chosen to match the Ernst angle independently for participant and sequence. The dataset with a 1s TR dataset was used to create the visual cortex regions of interest used in the following sections, using the contrast to non-zero faces, and adjusting the threshold until voxels that were in early visual cortex were isolated. These maps were then resampled for the more rapid acquisitions. While it would be preferable to use a separate localizer, as this circular selection does lead to a bias in favor of the slowest sampled dataset (1000ms), this bias cannot explain our primary finding of benefits in faster sampling (see below).

**Table 1.** Sequence parameters used for the empirical demonstration data. TR: Repetition Time; TE: Echo Time.

| TR (ms) | Multiband | In-Plane<br>Acceleration<br>(GRAPPA) | Partial<br>Fourier | Spatial<br>Resolution<br>(mm) | TE (ms) | Bandwidth<br>(Hz) | Number of<br>Slices |
| --- | --- | --- | --- | --- | --- | --- | --- |
| **1000** | 5 | 2 | 7/8ths | 1.6x1.6x1.6 | 22.0 | 1923 | 85 |
| **590** | 5 | 2 | 7/8ths | 2.0x2.0x2.0 | 15.6 | 2170 | 70 |
| **414** | 5 | 2 | 7/8ths | 2.5x2.5x2.5 | 16.4 | 2193 | 55 |
| **323** | 5 | 2 | 7/8ths | 3.0x3.0x3.0 | 14.2 | 2231 | 45 |

We processed the data in AFNI, performing slice time, motion and distortion correction and automated alignment to each participant’s anatomical image. Example images from each TR are shown in Figure 1, panel A. The images are similar, with the primary visible difference due to the voxel size. In order to confirm that the chosen sequence parameters successfully produced images with comparable SNR despite the changes in spatial and temporal resolutions, following (Dietrich et al., 2007), we calculated an image SNR metric by dividing the mean signal intensity in the gray matter by the standard deviation of the background (i.e. “noise”) signal.

Gray matter was defined using a mask of the cortical ribbon derived from FreeSurfer (Fischl and Dale, 2000) and resampled for the varying spatial dimensions of each of the acquisitions. Background signal was taken as the standard deviation within a sphere of 9mm radius, positioned outside of the head and regions of ghosting. Image SNR values were comparable across sequences, with a slight bias towards the 1000 ms TR, 1.6 mm isotropic voxel protocols (i.e. the HCP protocol). This is not surprising in light of the extensive effort (Glasser et al., 2016) devoted to optimizing the HCP acquisition pipeline (however note the largely overlapping error bars in Figure 1A).

Next, we computed the voxel-wise acquisition specific TSNR, defined as the mean of the signal in each run, after detrending with polynomials up to order 3, divided by the standard deviation. tSNR maps were warped into standard space independently for each sampling rate and averaged (Figure 1B). In contrast to image SNR, the tSNR shows increases as faster sampling is used. The highest tSNR is found for the most rapid acquisition and particularly in cortical areas. This is likely due to a number of factors, including reduced aliasing of physiological noise, better estimates of motion, better sampling of the HDR in addition to larger voxels (see below). Note that in the present dataset we chose to maintain identical acceleration factors across acquisition types in order to reduce the confound of SNR changes that can arise due to increased rates of acceleration (L. Chen et al., 2015). Furthermore, increases in tSNR are expected to saturate even as SNR continues to increase (Triantafyllou et al., 2011), implying that it is necessary to look beyond the tSNR metric in order to evaluate the benefits of increased temporal sampling.

In order to examine the statistical properties of the data, each run was scaled to have a mean of 100 and we then performed a GLM using the canonical HDR as implemented in AFNI (‘SPMG1’). The resulting beta and t-statistic maps were then warped into standard space and averaged for visualization purposes. Figure 1 panel C shows the mean beta values (only voxels with average t-statistic corresponding to p>0.01 are shown) from the contrast of all nonzero phase face stimuli versus baseline, conducted without (top) and with autocorrelation correction (bottom). Consistent with expectations, beta values show minimal differences between these two approaches, however, large differences are seen between different repetition times. In the 1000ms acquisition, the activation patterns show fine scale detail and larger amplitudes relative to the faster TRs, which also used larger voxel sizes, leading to partial volume effects. Though there is a loss of spatial precision due to the design of this experiment, there are benefits of rapid temporal sampling which will be discussed in further detail in a subsequent section.

In contrast to the lack of differences between betas, Figure 1, panel D shows the t-statistics maps from the two respective models. While a TR of 1000ms has very similar t-statistics between the two different approaches, larger differences are seen for the faster TRs. Two directly related effects are observed. First, the activation extent is reduced, as fewer voxels survive the arbitrary p<0.01 threshold. Second, the magnitude of the t-statistic is reduced, particularly in the case of the 323ms dataset, when autocorrelation correction is used. The peak magnitude of the t-statistics are reduced as the autocorrelation correction reduces the inflated t-statistics.

It should be noted though that, in light of the concomitant changes in temporal sampling rates and spatial resolutions across sequences, the observed tSNR gains are due, in part, to the larger voxel sizes, which increase as TRs get faster. In order to examine this possibility, the 1000ms data were smoothed, in Fourier space, to produce 3mm-equivilent data and resampled. These simulated 1000ms, 3mm data were then subjected to identical processing and analysis steps. tSNR was calculated as before showing that much, but not all, of the tSNR gains seemed to be related to voxels volumes (Figure 2A). Importantly, the activation extent (Figure 2B,C) and the peak of the t-statistic values remained significantly larger for the rapidly sampled data (Figure 2C), even when accounting for temporal autocorrelation. These findings suggest that the larger voxels do contribute to the higher tSNR measures, however the improvements above and beyond the smoothing effects support the conclusion that at least part of the observed improvements are related to faster sampling. As mentioned previously, these improvements could be related, amongst other things, to better sampling of the HDR (thus capturing more information) or reducing the relative impact of sudden subject movements.

**Figure 2.**
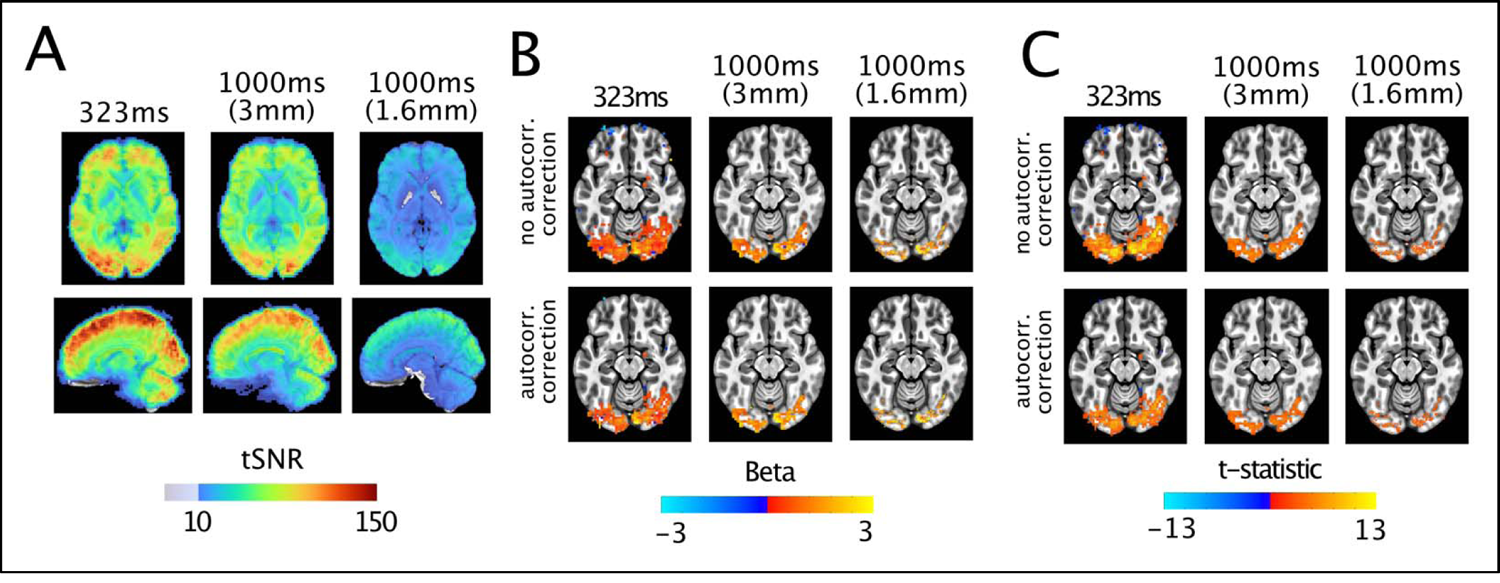
Characteristics of simulated 3 millimeter, 1000ms data. A) tSNR measures, average over subjects. The simulated 3mm data show gains in tSNR, but remain below the peak obtained with a sampling rate of 323ms. B) Activation amplitudes (betas) for the nonzero phase face stimuli compared to baseline. The impact of autocorrelation on betas is minimal, with the simulated data showing a larger, but still relatively smaller extent of activation. D) T-statistics for the same contrast. The 1000ms data is again little changed by simulating 3mm data acquisition and the peak t-statistics remain larger in the data sampled at 323ms. For visualization purposes, the beta and t-statistic maps use a t threshold of p<0.01 from the average t-statistics of original 1000 and 323ms data are reproduced from Figure 1.

Collectively these results highlight how, in an acquisition strategy where image SNR is matched across different protocols, temporal SNR can still increase with faster sampling rates and, additionally, reasonable patterns of activation can be generated for each acquisition. These findings also show the wide range of spatial-temporal scales available to investigators. One could also imagine reducing the coverage in order to achieve comparable temporal resolutions for a given spatial resolution, thereby gaining benefits of both increased spatial and temporal sampling.

### Denoising approaches can benefit from faster sampling

High frequency physiological noise arises primarily from the oscillatory nature of cardiac pulsations, which are typically around 1 Hz (Chen et al., 2019). In addition, there are slower physiological noise sources which include variation in the heart rate (Chang et al., 2009), respiration (Birn et al., 2008) and linked factors such as arousal (Chang et al., 2016). While traditional fMRI sometimes samples fast enough to capture variance driven by processes such as respiration (TR of ∼1.5s), cardiac signals require substantially faster (<500 msec) acquisitions. Under the slow TR fMRI framework these signals do not disappear, but are instead aliased into lower frequencies, corrupting the portion of the spectrum in which task responses or resting state fluctuations appear. With sufficiently fast sampling rates, it is often believed that it is possible to remove the unaliased physiological signals via filtering, however, this ignores higher order harmonics (Chen et al., 2019). Encouragingly, modeling methods that use cardiac and respiratory recordings can be used (Glover et al., 2000; Kasper et al., 2017) to generate regressors that capture and remove physiological noise. A limitation of this approach is that researchers often do not collect physiological recordings or are unable to acquire sufficiently clean recordings. Of course, if the physiological signals are corrupting the data, perhaps it is possible to derive the waveforms in a data driven manner. This idea was present in the early days of fMRI, with work using the phase information present at the center of k-space (‘frequency’ space, i.e. the Fourier transform of the image) to estimate respiratory and cardiac signals (Le and Hu, 1996). Recent work has conceptually built on this approach, reconstructing these physiological signals in a data driven manner when TRs are sufficiently short. Using an approach labeled HRAN, a statistical model of harmonic regression with autoregressive noise, Agrawal and colleagues show that with fast enough sampling, estimates of cardiac and respiratory noise can be produced from the fMRI data alone (Agrawal et al., 2020). These estimates performed as well as physiological noise regressors from RETROICOR and did not require separate physiological noise recordings.

Another approach involves examining time lagged correlations of the cardiac trace, as measured from pulse oximetry, in order to map its time lag across the brain (Tong et al., 2014). When sampling is sufficiently fast, it can be seen that the cardiac signal has effects across the whole brain, though the timing varies – an observation that requires rapid sampling to examine. This method can also be used to examine the low frequency fluctuations that are present in fMRI data, and again requires rapid sampling to reduce the aliasing of cardiac and respiratory signals into the lower frequencies of interest (Hocke et al., 2016). Though some harmonics may remain aliased even at fast sampling rates, there is a clear benefit of faster sampling in uncovering and potentially removing these confounding effects. At a minimum, fast sampling reduces the amount to which high frequency physiological noise components are aliased into the lower frequencies which are typically of interest for fMRI.

In addition, structured noise (which include noise from sources such as bulk motion) removal processes do appear to benefit from shorter TRs. With fast sampling, independent component analysis (ICA) methods appear to have more success in removing physiological noise and motion (Boubela et al., 2014; Griffanti et al., 2014), which can be applied in both task and resting state analyses. Multi-echo fMRI, a method that captures multiple “exposures” of the BOLD contrast in a single volume acquisition, can be combined with ICA methods for denoising (Kundu et al., 2012). With multi-echo fMRI contrast, denoising can be done based on whether there is a dependence of the signal on the echo time and then classifying signals as either BOLD or noise-like effects (Kundu et al., 2017, 2012). Multi-echo denoising used with ICA can also benefit from faster sampling rates (Boyacioğlu et al., 2015; Olafsson et al., 2015). Other denoising methods, such as GLMDenoise (Kay et al., 2013), which uses cross validation across multiple runs and derives noise regressors using principle component analysis (PCA) after determining a subject specific HDR, is also expected to perform better with faster sampling. Broadly speaking, any method which attempts to partition out insufficiently sampled temporal components of the signal is expected to benefit from faster sampling.

In addition to structured noise sources as discussed above, thermal noise is also a concern in fMRI. Thermal or system noise is Gaussian in nature and its relative contribution to the image increases with finer spatial sampling (Triantafyllou et al., 2011, 2005) or with the use of accelerated techniques (Todd et al., 2017). In rapid temporal sampling, lower flip angles are used, reducing the (arbitrary) signal intensity (in a given volume of tissue) of the image and its ratio to thermal noise. Unlike physiological noise sources, which are more detectable with faster sampling, there is no such benefit for thermal noise. Retrospectively, the thermal noise contribution in fMRI has typically been suppressed using spatial smoothing, which necessarily reduces spatial precision and can erroneously shift peak activation (Geissler et al., 2005; Jo et al., 2008; White et al., 2001). In high-speed fMRI, this may be even less desirable, as spatial resolution sacrifices have already been made to achieve the desired temporal resolution. Recent developments have led to methods that use low-rank patch based PCA methods to remove, in part, noise that cannot be distinguished from Gaussian (i.e. thermal noise) in the fMRI timeseries (Vizioli et al., 2020b). This method, known as NORDIC, can lead to increased t-statistics without the image blurring that is associated with spatial smoothing (Vizioli et al., 2020b). Here we show, using the previously introduce 323 millisecond dataset, that the NORDIC method is able to suppress thermal noise in rapidly sampled data. The NORDIC method was applied to these data before any processing. Subsequently, identical processing methods were applied to the images (see Empirical effects of faster sampling). This led to substantially less variance in the timeseries, and additionally, following the suppression of thermal noise, the contribution of physiological noise sources is clearer (Figure 3). The top left shows this for a single subject single voxel case in which the reduction of noise in the timeseries is clear, while the overall structure remains intact. This effect is apparent in the frequency domain after a fast Fourier transform (FFT), in which the respiratory (at ∼0.3Hz) and cardiac (∼1Hz) frequency peaks can be clearly distinguished after NORDIC but are buried in the high levels of thermal noise in the original data. These effects are also clear in the data after meaning over 419 voxels in the visual cortex ROI. While the mean timeseries are nearly identical, the FFT plot shows a clear reduction in power across a broad band of frequencies. As faster sampling become the norm, there is a clear need for methods that are able to suppress the relatively larger contribution of thermal noise. This is particularly true for the higher signal frequencies that are of interest and often contain physiological information that may be difficult to resolve due to thermal noise.

**Figure 3.**
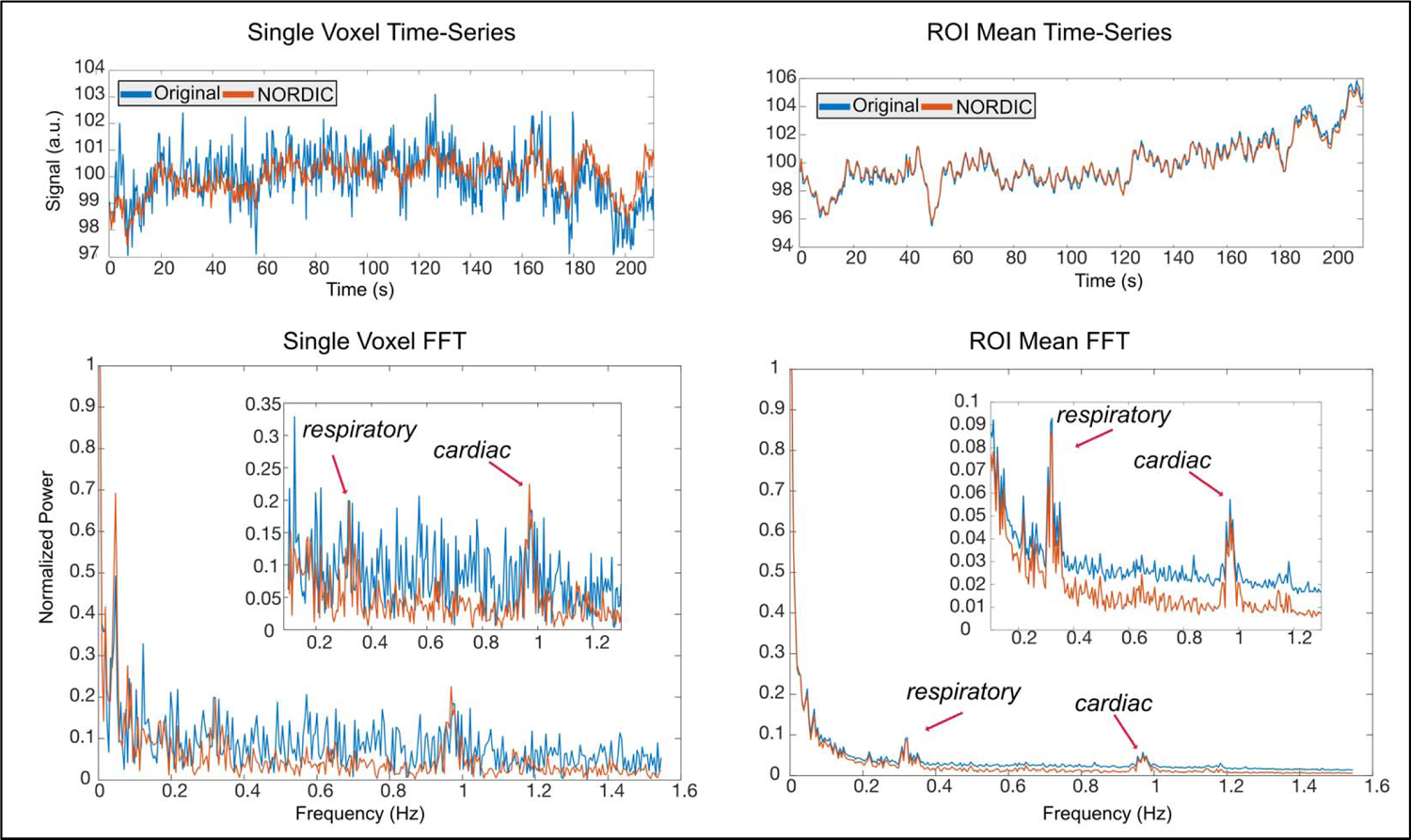
Timeseries and FFT plots show effects of NORDIC processing for 323ms data. Top Left. The timeseries for a single voxel is shown for the original (blue) and NORDIC-processed (Orange) data. The original data shows a larger amount of spurious noise, which is suppressed following NORDIC processing. Bottom Left. The FFT plots for this single voxel show that the effects of NORDIC are present across the entire power spectrum, though the respiratory and cardiac peaks at 0.3Hz and 1Hz remain in the data and are more easily seen. Top Right. Using the mean within the 419 voxels of the visual cortex ROI, there is little difference in the timeseries before and after NORDIC processing. Bottom Right. The FFT plot again shows a large suppression of noise, with physiological noise sources left intact.

Collectively, these findings show that there is a synergistic effect of faster sampling and denoising. These methods, whether they are tuned to remove physiological noise, other structured noise, or even thermal noise, appear to benefit from the increased dimensionality and reduced temporal aliasing achieved with fast fMRI acquisitions. This in turn produces better estimates of the hemodynamic response, higher t-statistics, and/or better parameter estimates (Bhandari et al., 2020). For example, going from a sampling rate of 1000ms to 323ms reduced the variance in single trial beta estimates, yielding a 57% (±6.2%) reduction in the standard deviation across trials for the 40% phase condition in the visual cortex ROI. The subsequent applications of NORDIC in the 323ms dataset, led to a further reduction of 9.5% (±6%) in the standard deviation of single trials, consistent with the effects described in prior work (Vizioli et al., 2020b). More work is required to sufficiently tease apart the benefits and downsides of faster imaging, as it is plausible that optimal analyses strategies will lead to more substantial analytical gains that have yet to be realized using methods that were developed for much slower sampling rates.

### Sampling rate interactions with common preprocessing steps

Estimating and correcting for motion is a critical step in the fMRI analysis pipeline. Faster sampling can allow for a more precise snapshot of motion, depending on the time scale of the motion relative to the TR. Volumes are aligned to one another, typically using a least-squares rigid body approach (i.e., 6 degrees of freedom), which reduces but does not entirely remove the effects of movement. Faster imaging could lead to more accurate motion estimates since motion might be captured instead of aliased, producing better outcomes after motion correction. However, there is a concern in that motion parameters in fast sampling regimes suggest that participants are constantly moving, with previously unnoticed motion occurring in the phase-encode direction of the EPI scan (Power et al., 2019). These effects have been studied, finding that they are due to a number of factors (Fair et al., 2020; Power et al., 2019). Namely, respiration is driving motion effects that are both “real”, in that the head is moving due to breathing, as well as apparent motion of the head resulting from magnetic field shifts due to movement of the chest during breathing (Power et al., 2019; Raj et al., 2001). In order to appropriately correct for motion, it may be necessary to notch filter the motion estimates in the respiratory band (Fair et al., 2020). Alternatively, it may be possible to correct for the effects of respiration at the time of acquisition either with dynamic shims (corrections for magnetic field distortion) applied during data acquisition (Stockmann and Wald, 2018; van Gelderen et al., 2007) or in a data-driven manner (Durand et al., 2001).

One other consideration is the impact of the slice timing correction. In typical 2D acquisitions, each slice (or slice group for multiband) is acquired at a unique point in time, however, the entire volume is considered a single timepoint by default. Particularly in event related studies (Sladky et al., 2011) correcting for within volume differences in timing improves accuracy and can be critical. The process of doing this does incur some penalty due to the necessity of interpolating the unmeasured timepoints and recent work has shown that slice-timing prior to motion correction can yield incorrect motion estimates, reducing the magnitude of sudden movement in particular (Power et al., 2017). With faster temporal sampling, the need for slice timing corrections, and thus interpolation, is reduced (Sahib et al., 2016; Sladky et al., 2011) or even eliminated.

### Neural Dynamics

#### Relationship between sampling and task effects

Faster sampling can also improve task separation. While amplitude has long been used as the key separator for task activation within the general linear modeling (GLM) framework, two slightly shifted HDRs can have identical amplitudes, despite very different response profiles (Lindquist et al., 2009).

Unbiased deconvolution models, such as those using finite impulse response (FIR) approaches, which directly estimate the HDR, have allowed for detailed investigations into the temporal dynamics of the BOLD signal. Under the FIR framework, a researcher selects a temporal window, or the number of TRs they expect the stimulus response to last, in which to estimate effects (Dale, 1999; Glover, 1999; Ollinger et al., 2001). The design matrix for the GLM then consists of a series of delta functions corresponding to the length of the time window multiplied by the number of unique stimulus conditions. This flexible and unbiased (i.e., no assumption of shape is made) approach produces a series of betas for each stimulus condition after the GLM, corresponding to the temporal estimates of the HDR. This approach can be used to deconvolve stimulus responses without making assumptions about specific shapes (Glover, 1999; Lewis et al., 2018). While FIR computations avoid mismodeling and mistaking temporal differences as amplitude effects, they do have the potential to overfit noise (Kay et al., 2008; Lindquist et al., 2009). Thus, their usage requires careful consideration of data quality and the chosen temporal window.

#### Empirical demonstration of the benefit of faster sampling: 1. Identifying task differences over time

To demonstrate the benefit of rapid temporal sampling within the FIR framework, which can be used to uncover differences between tasks that may be overlooked with slower sampling rates, we examined the multiple TR dataset introduced previously. In contrast to the GLM using the canonical HDR, we conducted a GLM using the finite impulse response (FIR) approach with a window of approximately 12 seconds in response to the 5 stimulus conditions (0 to 40% phase) for the separate face detection and fixation detection runs. We then extracted the FIR responses from a ROI that consisted of the primary visual cortex and compared all voxels FIR responses across all phases for the face detection against the fixation task (Figure 4).

**Figure 4.**
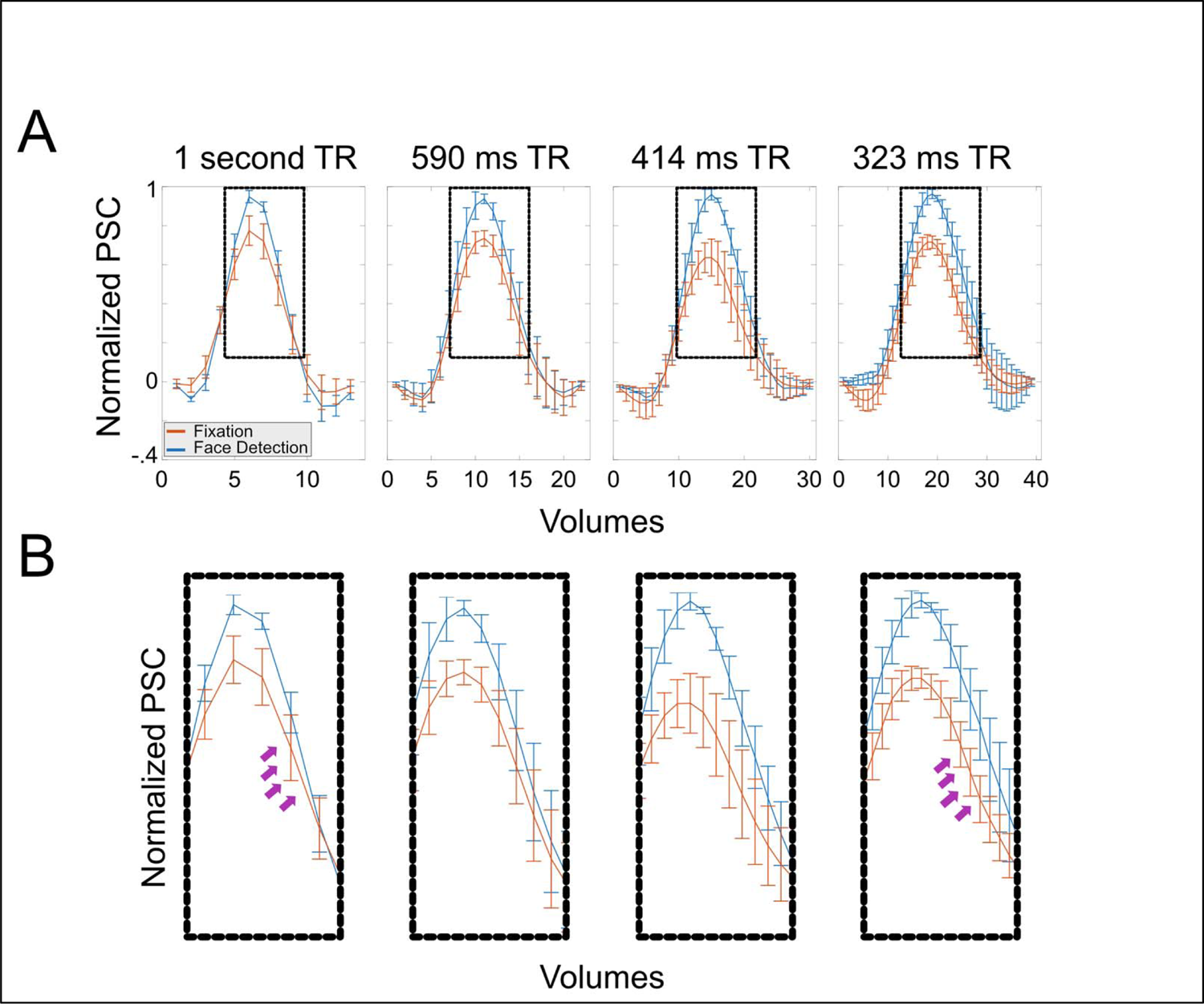
Clearer task differences at faster sampling rates. A) Panel A shows the average FIR (mean across subjects and phase coherences) for all resolutions elicited in the primary visual cortex during fixation and face detection task. Error bars portrays standard errors across subjects B) Zoomed in version of the dotted boxes in panel A. Note: 1. how latency differences across tasks become more prominent as TR decreases, particularly between 1000 and 323ms TR (arrows); 2. How amplitude difference across tasks also become more pronounced as TR decreases.

Our data indicate that as sampling rates increase, a more precise characterization of the HDR leads to a more precise quantification of temporal differences across tasks. Consider the 1 second TR acquisition (Figure 4, First column) in which 2 timepoints corresponding to peak of the curves can be seen as differentially responding to the 2 tasks (shown in different colors - column 1, lower; non overlapping error bars). For the 1 second TR, the HDR elicited by the face detection and fixation task are nearly perfectly overlapping, indicating no temporal differences across tasks. As we examine time courses with faster sampling, not only do we see more timepoints differentially responding to the 2 tasks (culminating in in 12 or more timepoints at a sampling rate of 0.323); but also, importantly, we begin to appreciate differences in the temporal structure of the HDRs (Column 4). More specifically, the response elicited by the fixation task drops off faster than that elicited by the face detection task (clearly seen for the .414 s and the .323 s TRs). Though preliminary, these findings highlight one of the benefits of rapid sampling in increasing the discriminability of task effects on the basis of HDRs estimate and show how this effect varies over the duration of the response.

Moreover, as we sample data faster more prominent temporal structure can be appreciated. For example, for the fixation task, the amplitude of the initial dip increases as a function or temporal resolution, ranging from an average amplitude of approximately −.01±0.036 (mean ± standard error) for the 1 second TR to −0.093±0.047 for the 0.323 second TR (Figure 5, Top). Here we are showing an improved ability to detecting early temporal structure (i.e., initial dip) as TR decreases, however, these datasets varied across several parameters, including in their spatial resolution and tSNR characteristics. This was done to ensure that image SNR was maximized and comparable across protocols (see Figure 1). To further compare temporal characteristics across temporal resolutions only (that is, keeping all other parameters constant), we simulated 3 slower TR datasets by downsampling the 323ms data. These were produced by averaging 2, 3 or 4 neighboring timepoints to produce simulated data with effective TRs of 646, 969 and 1292ms. Identical processing method and analyses were conducted to determine how this change in sampling affected the observation of the initial dip (Figure 5, middle row). Despite being derived from the 323ms data, which possess a robust initial dip, we observed that downsampling the data dramatically reduced (646ms data) or eliminated this effect (969 or 1292 data). These findings suggest that sampling rates are driving these effects, and it is not an artifact of the changing voxel size or tSNR characteristics.

**Figure 5.**
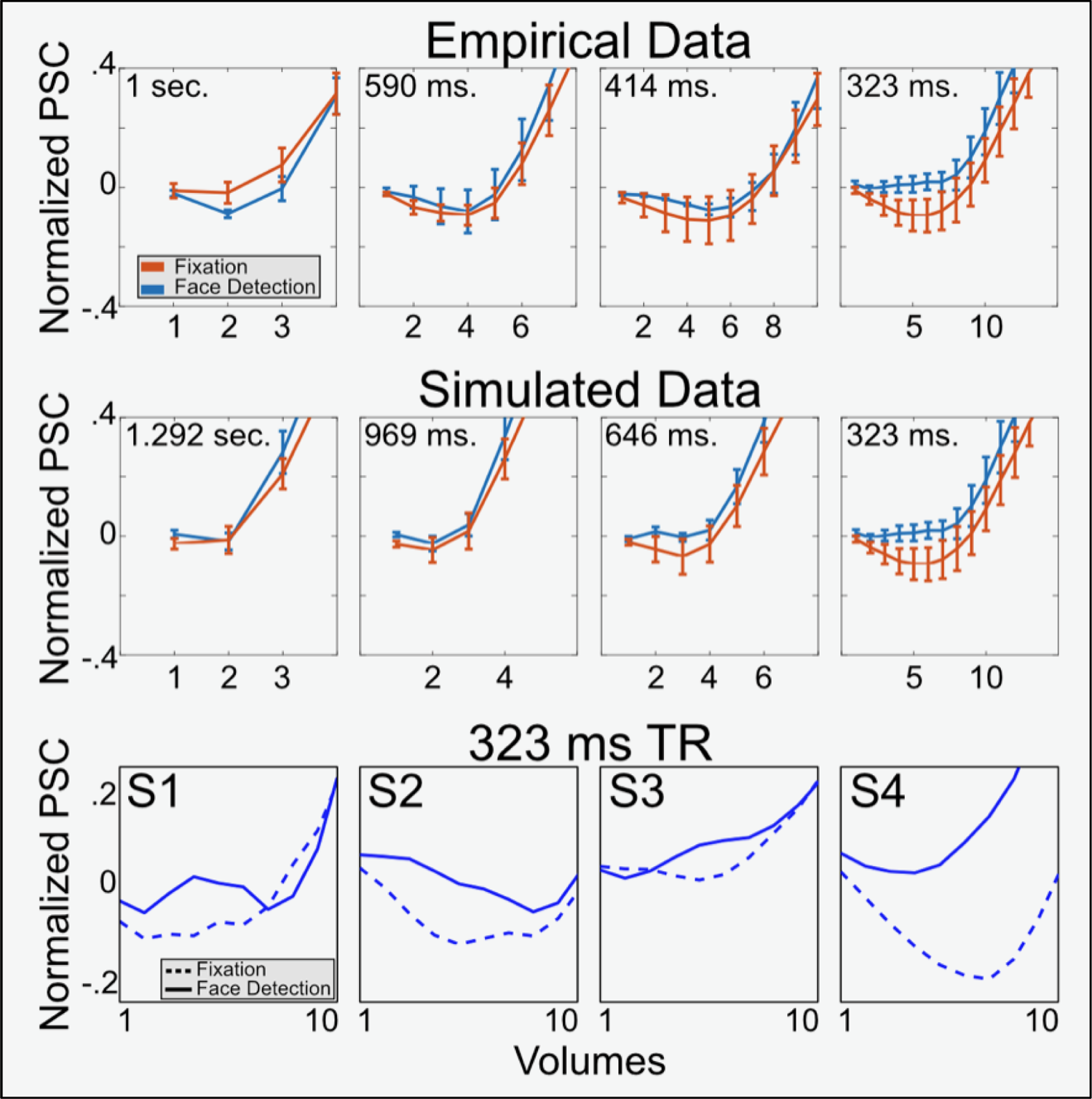
Clearer emergence of initial dip with sampling rate increases. The top two panels illustrate how the initial dip becomes more pronounced as TR decreases. Specifically, task differences in the initial dip only emerge at the fastest sampling rate (i.e., 323ms TR). Both top panels zoom in the initial dip for the average FIRs across subjects and conditions elicited by both tasks in the visual cortex ROI, with error bars representing standard error across subjects. The t shows the initial responses in the empirical data for all resolutions (as portrayed in Figure 3). The middle panel i shows downsampled data, simulating different TRs. Note the striking correspondence between empirical and si p panel stead ulated data. The bottom panel instead shows the first 10 volumes (i.e., 0 to 3.23 seconds after stimulus onset) for each single subject and for both tasks, only for the 323 ms TR data (i.e., the fastest temporal sampling), which the data set s owing the earlies differences across tasks. Note how all subjects show an increased negativity for fixation compared to face detection.

More importantly, also as a function of temporal resolution, our analyses indicated early emergence of attentional differences during the period of the initial dip. We found that, for the fastest resolution only, all subjects showed task differences emerged as early as 5 TRs (i.e., ∼1.62 secs) after stimulus onset (Figure 5, Bottom row), with larger responses (i.e., more negative) for the fixation (amplitude: −0.095±0.053 at TR 5) than the face detection task (amplitude: 0.01±0.027 at TR 6). These timings are consistent with those reported for the initial dip (Hu et al., 1997; Menon et al., 1995) and suggest that sufficient temporal sampling is required to detect such rapid and (across all subjects – see Figure 5) small attentional effects.

The initial dip, which was noted to begin within the first second of a BOLD response (Hu et al., 1997; Menon et al., 1995), has been difficult to confirm (Buxton, 2001), with a number of studies not being able to identify this temporal component, despite ample averaging (Fransson et al., 1999, 1998). A number of variables have in fact been attributed to the presence or absence of the initial dip, including field strength, task timing, stimulus type, echo time and spatial and temporal dimensions of the sampled data (Hu and Yacoub, 2012; Watanabe et al., 2013) and it is reasonable to expect that these causes can interact. It is, of course plausible that subtle differences between the task conditions in early visual cortex are giving rise to temporal differences in the initial dip and more generally across the HDR. As such, with typical acquisition strategies this effect is missed (Figure 4, first column), leading to incorrect assumptions about task demands or task effects. Here we report that attentional differences related to task can also be visible in the initial dip, which has been suggested to reflect more localized neuronal responses (Hu et al., 1997; Menon et al., 1995; Yacoub and Hu, 2001). While this observation further supports the value of temporal information in fMRI, the size, transient nature, individual differences and magnitude of these effect further advocate the need to continue developing tailored tools to extract this information.

As the data presented here are meant to provide insights into the temporal dimension of BOLD responses, a thorough neuroscientific interpretation of these results is beyond the scope of this work. However, the differences in the amplitude of the initial dip across tasks could be related to response inhibition during the fixation task, where faces are considered as distractors and must be suppressed.

Generally, we report both early (i.e. 5 TR after stimulus onset) and late (after the HDR peak) latency differences become clearer as temporal resolution increases. Albeit preliminary, due to the small N, these promising results suggest that, if fine grained temporal information is available, faster temporal sampling may uncover additional temporal structure or stimuli related effects that were previously overlooked. Of course, finding such effects may require specific analysis strategies tailored for voxel wise HDRs and single trial estimates of responses.

### Temporal dynamics in fMRI

Historically, rapid (i.e. sub-second) temporal dynamics in fMRI have been for the most part neglected due to the inability of acquiring rapidly sampled data in conjunction with both reasonable spatial resolution and/or whole or near whole brain coverage. More recently, optimizations in both software and hardware have allowed the recording of functional images with unprecedented spatial and temporal resolutions, while maintaining reasonable SNR and CNR. Consequently, more and more attention has been devoted to statistical analyses of TR-to-TR fluctuations in the sub-second regime.

For example, in the realm of spatial analyses, considerable attention has been given to spatial clustering (Cox et al., 2017; Eklund et al., 2016; Hayasaka and Nichols, 2003; Nichols, 2012), including using Monte Carlo methods (Cox et al., 2017). Advanced multivariate methods have also been developed to account for spatial patterns of activity (Haxby et al., 2014, 2001), including using search light approaches (Kriegeskorte et al., 2006). Notably, these spatially oriented methods can also benefit from temporal detail. For example, Vu and colleagues demonstrated that accurate word timing, made possible by fast TR fMRI data, improved the performance of multivoxel pattern analyses (MVPA) (Vu et al., 2016).

Recently, however, there has been some development in methods which directly consider temporal dynamics. For example, when considering deconvolution approaches, it is necessary to correctly deal with the multiple levels of statistical interdependence. A voxel-wise, linear mixed modeling approach was developed to examine both where and when statistical differences occur in complete HDRs (G. Chen et al., 2015). The benefit of this method is that it allows the statistical examination of a full curve, rather than summarizing the multiple parameter estimates via averaging or area-under-the-curve calculations. This becomes increasingly important as sampling rates and thus the number of timepoints estimated in the deconvolution increase.

Analogous to multivariate pattern analysis for spatial maps, a recently developed method considers the temporal corollary of this, termed temporal MVPA (tMVPA) (Vizioli et al., 2018). tMVPA uses single trial response time courses or single run FIR estimates to compute single trial Representation Dissimilarity Matrices (RDM) independently per condition within a given ROI. To infer statistically significant differences across time courses, tMVPA builds on a sliding window approach (extensively tested on real and synthetic data elsewhere – Vizioli et al., 2018) that allows for the precise identification of the temporal window of an effect and whether this encompasses only a few time points or is sustained over a larger time window. Multivariate analyses methods have been shown to have increased sensitivity for the analyses of spatial maps (Kriegeskorte and Bandettini, 2007; Vizioli et al., 2020a) in fMRI. There is evidence that this is also the case for the temporal domain, with prior work finding earlier identification of statistically significant task differences on both real (Ramon et al., 2015) and synthetically simulated data (Vizioli et al., 2018).

### Partial windowing of temporal effects

Partial volume effects (i.e. when the prescribed voxel size is not small enough to capture only a single tissue) are often discussed in the fMRI literature. This describes a phenomenon in which a single voxel can have contributions from multiple tissue types, or from two nominally independent cortical layers (Siero et al., 2013). In the temporal domain, a parallel manifestation of this is aliasing of oscillatory signals due to sampling slower than the Nyquist frequency. This is typically considered in the context of physiological noise regressors, such as cardiac pulsation (Chen et al., 2019). For example, cardiac signals, which tend to range between 0.8 to 1.5Hz require rapid sampling rates in order to avoid aliasing, even faster than what is used for increasingly common sub-second TRs. While rapid sampling does improve the fidelity of these signals and aid in their removal (Chen et al., 2019; Tong et al., 2014) we wish to highlight additional considerations beyond undesirable physiological noise.

Unlike periodic physiological signals, there is often little consideration of any loss of task relevant BOLD signal information due to insufficient temporal sampling. However, given the rich dynamics of the BOLD signal and the short delay between a stimulus and a potential initial dip or initial rise in signal intensity, there is a risk that insufficient sampling rates can lead to “partial windowing” of temporal signals of interest. This effect can manifest due to their short duration and magnitude, as is the case for the initial dip. To some extent, this can be mitigated by jittering the stimulus onset with respect to TR (Amaro and Barker, 2006; Dale, 1999; Miezin et al., 2000; Watanabe et al., 2013), but this requires jittering with sufficient time steps in order to capture all temporal aspects of interest, i.e. on the order of 500 ms or shorter. Considering the duration of a conventional HDR (on the order of 20s), this would require multiple stimulus presentations to capture a single “full” HDR, producing overly long scan times or reduce repeats of the stimulus. This is also not to say that faster acquisition times cannot take advantage of jitter, however, they would require substantially fewer variants of the stimulus timing to sample all of the temporal characteristics of interest, thereby reducing “partial windowing” effects. If sampling is sufficiently fast, then the effect of partial windowing is reduced even further, producing estimates of the HDR that capture fast temporal dynamics (Figure 5, above).

### High frequency and dynamic correlations in resting state

If the effects of interest are oscillatory in nature, then jittering is no longer a reasonable option and instead rapid acquisitions may be more suited for observing high frequency effects. For example, in resting state fMRI, the primary frequencies of interest appear in low frequency bands, between 0.1 and 0.01Hz. For these frequencies, faster sampling primarily serves to reduce the impact of physiological noise, as discussed previously. Such low frequency bands are easily sampled with classic 2 to 3 second TRs and would, by necessity, contain substantially higher relative power when compared to high frequency fluctuations. Recently, however, with the advent of faster sampling, there have been observations of resting state correlations at frequencies greater than 1Hz (Gohel and Biswal, 2014; Smith-Collins et al., 2015). Though this work is still ongoing, there are methodological concerns as denoising procedures make it possible to reintroduce noise under the linear regression framework (Lindquist et al., 2019) and even produce spurious or enhanced high frequency correlations (Chen et al., 2016).

One argument against high frequency correlations concerns the plausibility of their origin. Based on conventional models of the BOLD response, it should be incredibly difficult to detect neuronally driven oscillations at such high frequencies (Chen et al., 2016), however it has been suggested that through suitable task designs it is possible to derive information from effects in the 10s of millisecond range (Ogawa et al., 2000). This is ultimately an empirical question that can be evaluated using oscillating stimulus designs with sufficiently high frequencies. Unlike relative timing within regions or between separate tasks and stimuli, oscillatory or incredibly rapid stimulus presentation is much more difficult to detect. This is due to the plateauing nature of the BOLD response, converting these rapid oscillations into small fluctuations. Recent work has uncovered that with sufficiently fast sampling, fMRI can capture rapid oscillations in visual stimuli up to 0.75 or potentially 1Hz (Lewis et al., 2016) and other have found that neurovascular coupling appears to occur on shorter timescales than were typically thought (Siero et al., 2011; Silva and Koretsky, 2002). How these findings have led to a reconsideration of the dynamics of the HDR and its underlying biological principles are considered in a separate manuscript in the current issue (Polimeni, and Lewis, 2021).

An alternative extension to resting state analyses, known as dynamic functional connectivity (dFC), consider the change in the correlation pattern over time, often within sliding windows containing 30 to 60s worth of samples (Hutchison et al., 2013; Preti et al., 2017). These studies seek to explore how brain states change, in contrast to static functional connectivity which captures a single measure of correlations across the brain. This approach has been used, for example, to link behavioral measures of executive performance with the changes in network state (Braun et al., 2015) or link changes in the default mode network while listening to narratives with memory retrieval (Simony et al., 2016). One recurring question for a windowed approach concerns the window length. With a TR of 2 seconds, there is often a difficulty in selecting an adequate window length that balances statistical power concerns against capturing each dynamic state (Hutchison et al., 2013). Here the statistical gains of rapid sampling can be directly linked to the benefit of uncovering neural dynamics, as there is evidence that faster sampling provides for gains in measuring significant functional connectivity changes within shorter and shorter windows (Sahib et al., 2018) or capturing network effects that are missed in static analyses of clinical populations (Zhang et al., 2018). Though resting state emerged as a methodology focused on slow (0.1 to 0.01Hz) oscillations for which typical 2s TRs are more than adequate, these findings suggest that there are substantial benefits and novel neuroscientific findings waiting to be uncovered with fast fMRI.

### Combining fMRI and EEG/MEG

Tools such as electroencephalography (EEG) and magnetoencephalography (MEG) offer high temporal resolution for human imaging but suffer from limited spatial precision. By combining these techniques with fMRI, concurrently in the case of EEG, or consecutively for MEG, it is possible to combine the precise spatial information with the rich temporal signals from electrophysiology.

In line with the goal of clinical utility for fMRI, there is ongoing research in epilepsy, as concomitant spatial and temporal precision could prove extraordinarily useful for presurgical planning, for example. Even with TRs of 3000ms early work showed that these methods were not only feasible but could potentially augment the limited spatial precision of EEG, though effectiveness varied over the sample (Salek-Haddadi et al., 2006). However, there is evidence that faster fMRI methods may have superior performance (Jacobs et al., 2014) and clinical promise. Using a rapid functional imaging technique termed magnetic resonance encephalography (MREG) (Assländer et al., 2013; Zahneisen et al., 2012), Jäger and colleagues were able to obtain whole brain data with 100ms temporal sampling and reasonable (4 to 5mm) voxel resolutions. In patients with epilepsy, they found increased support for positive BOLD responses co-occurring with epileptic spikes, though further work is required to better understand the link between the highly variable spikes and the resulting HDR (Jäger et al., 2015).

Furthermore, in the case of simultaneous EEG/fMRI, the temporal information available from EEG can be used to supplement and better understand the co-occurring fMRI signal (Chang and Chen, 2021; Murta et al., 2015) or potentially confirm novel hemodynamic findings (Lewis et al., 2016). Studies combining EEG and fMRI have spanned the full range experimental paradigms routinely used in fMRI, from naturalistic stimuli (Whittingstall et al., 2010), to resting state (Deligianni et al., 2014; Goldman et al., 2002; Mayhew and Bagshaw, 2017; Meyer et al., 2013; Wirsich et al., 2020) and typical task based approaches (Hill et al., 2021; Mayhew and Bagshaw, 2017).

Using flickering gratings within task setting, for example, Lewis and colleagues found that HDRs in response to 0.5Hz and 0.2Hz oscillating stimuli were larger and more visible than expected by conventional HDR models. By recording simultaneous EEG, they were able to argue against these linear canonical models of the HDR, as the underlying neural activity (driving the EEG responses) did not differ strongly in magnitude (Lewis et al., 2016).

Though MEG cannot be acquired simultaneously with fMRI, it still offers promise for integration. Difficult fMRI latency investigations (See Hemodynamic Response Variability section) can be augmented with event timings derived from MEG to investigate and validate latency estimates derived from fMRI (Lin et al., 2013). Even in complex, putatively hierarchical stimulus processing, such as visual object recognition, combining MEG and fMRI experimental results (Cichy et al., 2016), yields millisecond resolution in the ventral visual stream processing cascade.

While there are several exciting research avenues associated with combining these methods, there are a number of limitations to consider. For example, in light of their different physiological origins, reconciling neuronal responses from hemodynamically driven BOLD effects and electrophysiological cortical signals can be a challenge (Rosa et al., 2010). In addition, EEG data acquired simultaneously with fMRI is corrupted with a number of artifacts, including those from the MR hardware, such as gradient effects (Allen et al., 2000; Yan et al., 2009) and those associated with the subject, such as the head motion or the ballistocardiograph artifact from the heartbeat (Allen et al., 1998; Debener et al., 2008; Masterton et al., 2007). Pushing simultaneous EEG/fMRI to higher field strengths (>=7 Tesla) increases the prominence of these artifacts and further degrades the BOLD signal (Jorge et al., 2015; Neuner et al., 2014).

Despite these challenges, these findings highlight the diverse benefits of combining the temporal resolution available with EEG or MEG with the spatial precision of fMRI – and show that despite the lagging BOLD response, there are benefits in accelerated fMRI when combined with these temporally precise methods.

### Relationship between spatial specificity and the temporal signal

While much of fMRI has pushed for higher and higher spatial resolutions, there is also a need to consider how temporal dynamics interact with spatial precision as a specific point in time gives rise to a non-stationary spatial activation map. For example, we know that the HDR changes across regions, but also within regions across cortical depths. It would be logical to assume that when temporal sampling is too low, accurate characterization of the model HDR (or of the empirical time courses themselves) would be unachievable. Consequently, model fit may be suboptimal, leading to misestimation of response amplitude parameters, thus producing inaccurate spatial maps. Here, we consider further implications for spatial resolution and specificity in the context of temporal information.

### Depth-dependent fMRI and spatiotemporal information

There is evidence that early and late phases of the HDR have greater spatial specificity relative to the central peak (Goodyear and Menon, 2001; Puckett et al., 2016; Shmuel et al., 2007). In line with this, it was proposed that fitting or analyzing just the early parts, such as the initial dip, could provide a method to gain spatial specificity (Hu et al., 1997; Menon et al., 1995; Yacoub and Hu, 2001). A similar possibility manifests in depth-dependent fMRI data. As venous blood drains through the cortical column, there is a slight delay between the peak of deep layers and the large veins on the pial surface (Petridou and Siero, 2019; Siero et al., 2011, 2015; Silva and Koretsky, 2002). This effect appears as differences in the HDR between layers, which must be accounted for in analyses. Using an identical model for the HDR across all layers will lead to mismodelling (Lindquist et al., 2009) and could produce spurious layer dependent effects if the temporal differences appear as amplitude changes. Alternatively, as a consequence of the same mismodelling, the true effects, which are often very small in magnitude, could be missed. HDR estimates must be carefully tailored for each layer in order to make depth dependent inferences about neural connections or activity.

Building on this idea, it is has been recently demonstrated that we can use this temporal information to separate out the earlier responses, which may be nominally more spatially precise, from the later responses (Kay et al., 2020). This method, termed temporal decomposition through manifold fitting (TDM), uses unbiased FIR estimates of voxel time courses to produce two regressors for each event, one early and one late. This method separates out these responses rather than accounting for timing differences, as is done with methods that use multiple basis functions. While this method works with data sampled at sampling rates of up to 2.2 seconds, the time courses of early and late responses are highly correlated, and higher temporal sampling would aid in separation and provide more data dimensionality for estimating early and late responses. More work is needed to consider how spatial effects interact with the temporal domain and what resolutions are required for an optimal experimental paradigm. It is likely that reduced field of view imaging will remain common, as whole brain imaging at both high spatial and temporal resolutions remains difficult.

### Line scanning as a preview of the future of fMRI

Accelerated fMRI can offer increases in precision and spatial specificity across larger sections of the cortex but remains relatively course on the scale of local neural activity. One technique, which takes the temporal and spatial focus to its extreme is termed line scanning (Albers et al., 2018; Yu et al., 2014). Typically performed in animal fMRI, line scanning offers unprecedented spatial <(.5 mm) and temporal (100 to 50ms) resolution with the limitation of collecting an extremely limited, one-dimensional portion of the cortex.

In rodents this has been used to interrogate the precise temporal differences between optogenetic stimulation relative to direct sensory stimulation (Albers et al., 2018), as well as confirming the layer specificity of fMRI responses (Yu et al., 2014). While these results were obtained under very specific acquisition paradigms, they support the quest for higher temporal and spatial precision, showing that this, under the right circumstances, can provide for neurobiological precision as well.

In humans, line-scanning presents new challenges, however, development has made initial explorations of this technique possible. Recent preliminary results in humans show that 0.2mm resolution with 100ms temporal precision is achievable, albeit over a single and selective portion of the human visual cortex (Morgan et al., 2020). By using a multi-echo sequence, this group was able to estimate the T2* parameter and its variability across cortical layers (Koopmans et al., 2011), obtaining increased functional precision. In addition, they find that this precise mapping technique recapitulates findings from electrophysiology, namely visual tuning properties (Morgan et al., 2020).

These results build upon the ideas of the previous sections, showing that even at the extremes of currently feasible temporal and spatial sampling there is rich information to be gained. Of course, the current field of view requirements prevent examining the interaction of local and long-range networks, limiting the type of research questions that can be asked for now. In addition, data collected at such high resolutions inherently has very low SNR, being dominated by thermal noise. These issues highlight the important of methods to reduce thermal noise, such as NORDIC (Vizioli et al., 2020b, see Denoising approaches section). In time, it is likely that the difficulties associated with these types of acquisitions and analyses will be further reduced, providing for a new frontier of spatiotemporal fMRI discoveries.

## Conclusion

Technical developments and the increasing availability of advanced pulse sequences and methods have made ultra-fast fMRI routinely possible. Here, we have tried to address a seemingly straightforward question – What are the benefits of increased temporal sampling for fMRI? In combining information from existing work and our own empirical demonstrations we have shown that the answer is complex, but ultimately encouraging. We have argued that the benefits are not just limited to statistical gains, but encompass a wide range of potential neuroscientific questions, including BOLD dynamics at rapid timescales. While we find consistent effects across subjects and convergence with emerging data, additional work, with larger sample sizes, will be required to confirm the specific timing of the effects reported. Nonetheless, the idea that faster temporal sampling can help extract fine grained temporal information, if it is available, is demonstrated by these data. Of course, additional considerations are needed regarding the challenges and difficulties associated with collecting, analyzing and interpreting rapidly sampled data. Nevertheless, the potential benefits are many: statistical power, better denoising, better spatial resolution and, importantly additional and more precise insights into neuronal temporal dynamics. Though space and time are intrinsically linked in fMRI, the relationship between accurate measures of temporal dynamics and spatial precision is often overlooked. More precise timing estimates can produce better models, which in turn will produce more accurate spatial maps. Collectively this diverse array of benefits suggests the continued work is worth the effort. With the combined efforts across hardware, pulse sequence developments, statistical understanding and methodological approaches the functional neuroimaging field is ready for a resurgence or perhaps a return, to the days of in depth, temporal investigations.

## Acknowledgments

Funding for this study was provided by National Institutes of Health Grants RF1 MH117015 (Ghose), RF1 MH116978 (Yacoub), P41 EB027061 (Ugurbil) and P30 NS076408 (Ugurbil).

## Bibliography

1. Agrawal, U., Brown, E.N., Lewis, L.D., 2020. Model-based physiological noise removal in fast fMRI. NeuroImage 205, 116231. https://doi.org/10.1016/j.neuroimage.2019.116231

2. Albers, F., Schmid, F., Wachsmuth, L., Faber, C., 2018. Line scanning fMRI reveals earlier onset of optogenetically evoked BOLD response in rat somatosensory cortex as compared to sensory stimulation. NeuroImage, Pushing the spatio-temporal limits of MRI and fMRI 164, 144–154. https://doi.org/10.1016/j.neuroimage.2016.12.059

3. Allen, P.J., Josephs, O., Turner, R., 2000. A Method for Removing Imaging Artifact from Continuous EEG Recorded during Functional MRI. NeuroImage 12, 230–239. https://doi.org/10.1006/nimg.2000.0599

4. Allen, P.J., Polizzi, G., Krakow, K., Fish, D.R., Lemieux, L., 1998. Identification of EEG Events in the MR Scanner: The Problem of Pulse Artifact and a Method for Its Subtraction. NeuroImage 8, 229–239. https://doi.org/10.1006/nimg.1998.0361

5. Amaro, E., Barker, G.J., 2006. Study design in fMRI: basic principles. Brain Cogn 60, 220–232. https://doi.org/10.1016/j.bandc.2005.11.009

6. Assländer, J., Zahneisen, B., Hugger, T., Reisert, M., Lee, H.-L., LeVan, P., Hennig, J., 2013. Single shot whole brain imaging using spherical stack of spirals trajectories. NeuroImage 73, 59–70. https://doi.org/10.1016/j.neuroimage.2013.01.065

7. Avossa, G., Shulman, G.L., Corbetta, M., Shulman, G.L., Iden-, M.C., 2003. Identification of Cerebral Networks by Classification of the Shape of BOLD Responses 360–371.

8. Bandettini, P.A., 2014. Neuronal or Hemodynamic? Grappling with the Functional MRI Signal. Brain Connectivity 4, 487–498. https://doi.org/10.1089/brain.2014.0288

9. Bandettini, P.A., Jesmanowicz, A., Wong, E.C., Hyde, J.S., 1993. Processing strategies for time-course data sets in functional MRI of the human brain. Magn Reson Med 30, 161–173. https://doi.org/10.1002/mrm.1910300204

10. Baumann, S., Griffiths, T.D., Rees, A., Hunter, D., Sun, L., Thiele, A., 2010. Characterisation of the BOLD response time course at different levels of the auditory pathway in non-human primates. NeuroImage 50, 1099–1108. https://doi.org/10.1016/j.neuroimage.2009.12.103

11. Bellgowan, P.S.F., Saad, Z.S., Bandettini, P.A., 2003. Understanding neural system dynamics through task modulation and measurement of functional MRI amplitude, latency, and width. PNAS 100, 1415–1419. https://doi.org/10.1073/pnas.0337747100

12. Bhandari, R., Kirilina, E., Caan, M., Suttrup, J., De Sanctis, T., De Angelis, L., Keysers, C., Gazzola, V., 2020. Does higher sampling rate (multiband + SENSE) improve group statistics - An example from social neuroscience block design at 3T. NeuroImage 213, 116731. https://doi.org/10.1016/j.neuroimage.2020.116731

13. Birn, R.M., Cox, R.W., Bandettini, P.A., 2002. Detection versus Estimation in Event-Related fMRI: Choosing the Optimal Stimulus Timing. NeuroImage 15, 252–264. https://doi.org/10.1006/nimg.2001.0964

14. Birn, R.M., Smith, M.A., Jones, T.B., Bandettini, P.A., 2008. The Respiration Response Function: The temporal dynamics of fMRI signal fluctuations related to changes in respiration. Neuroimage 40, 644–654. https://doi.org/10.1016/j.neuroimage.2007.11.059

15. Bollmann, S., Barth, M., 2020. New acquisition techniques and their prospects for the achievable resolution of fMRI. Progress in Neurobiology 101936. https://doi.org/10.1016/j.pneurobio.2020.101936

16. Bollmann, S., Puckett, A.M., Cunnington, R., Barth, M., 2018. Serial correlations in single-subject fMRI with sub-second TR. NeuroImage 166, 152–166. https://doi.org/10.1016/j.neuroimage.2017.10.043

17. Boubela, R.N., Kalcher, K., Nasel, C., Moser, E., 2014. Scanning fast and slow: Current limitations of 3 Tesla functional MRI and future potential. Frontiers in Physics 2, 1–8. https://doi.org/10.3389/fphy.2014.00001

18. Boyacioğlu, R., Schulz, J., Koopmans, P.J., Barth, M., Norris, D.G., 2015. Improved sensitivity and specificity for resting state and task fMRI with multiband multi-echo EPI compared to multi-echo EPI at 7T. NeuroImage 119, 352–361. https://doi.org/10.1016/j.neuroimage.2015.06.089

19. Boynton, G.M., Engel, S.A., Glover, G.H., Heeger, D.J., 1996. Linear Systems Analysis of Functional Magnetic Resonance Imaging in Human V1. J. Neurosci. 16, 4207–4221. https://doi.org/10.1523/JNEUROSCI.16-13-04207.1996

20. Boynton, G.M., Engel, S.A., Heeger, D.J., 2012. Linear Systems Analysis of the fMRI Signal. Neuroimage 62, 975–984. https://doi.org/10.1016/j.neuroimage.2012.01.082

21. Braun, U., Schäfer, A., Walter, H., Erk, S., Romanczuk-Seiferth, N., Haddad, L., Schweiger, J.I., Grimm, O., Heinz, A., Tost, H., Meyer-Lindenberg, A., Bassett, D.S., 2015. Dynamic reconfiguration of frontal brain networks during executive cognition in humans. PNAS 112, 11678–11683. https://doi.org/10.1073/pnas.1422487112

22. Breuer, F.A., Blaimer, M., Heidemann, R.M., Mueller, M.F., Griswold, M.A., Jakob, P.M., 2005. Controlled aliasing in parallel imaging results in higher acceleration (CAIPIRINHA) for multi-slice imaging. Magn Reson Med 53, 684–691. https://doi.org/10.1002/mrm.20401

23. Bullmore, E., Brammer, M., Williams, S.C., Rabe-Hesketh, S., Janot, N., David, A., Mellers, J., Howard, R., Sham, P., 1996. Statistical methods of estimation and inference for functional MR image analysis. Magn Reson Med 35, 261–277. https://doi.org/10.1002/mrm.1910350219

24. Buxton, R.B., 2001. The Elusive Initial Dip. NeuroImage 13, 953–958. https://doi.org/10.1006/nimg.2001.0814

25. Chang, C., Chen, J.E., 2021. Multimodal EEG-fMRI: advancing insight into large-scale human brain dynamics. Curr Opin Biomed Eng 18, 100279. https://doi.org/10.1016/j.cobme.2021.100279

26. Chang, C., Cunningham, J.P., Glover, G.H., 2009. Influence of heart rate on the BOLD signal: The cardiac response function. NeuroImage 44, 857–869. https://doi.org/10.1016/j.neuroimage.2008.09.029

27. Chang, C., Leopold, D.A., Schölvinck, M.L., Mandelkow, H., Picchioni, D., Liu, X., Ye, F.Q., Turchi, J.N., Duyn, J.H., 2016. Tracking brain arousal fluctuations with fMRI. PNAS 113, 4518–4523. https://doi.org/10.1073/pnas.1520613113

28. Chang, C., Thomason, M.E., Glover, G.H., 2008. Mapping and correction of vascular hemodynamic latency in the BOLD signal. Neuroimage 43, 90–102. https://doi.org/10.1016/j.neuroimage.2008.06.030

29. Chang, W.-T., Nummenmaa, A., Witzel, T., Ahveninen, J., Huang, S., Tsai, K.W.-K., Chu, Y.-H., Polimeni, J.R., Belliveau, J.W., Lin, F.-H., 2013. Whole-head rapid fMRI acquisition using echo-shifted magnetic resonance inverse imaging. NeuroImage 78, 325–338. https://doi.org/10.1016/j.neuroimage.2013.03.040

30. Chen, G., Saad, Z.S., Adleman, N.E., Leibenluft, E., Cox, R.W., 2015. Detecting the subtle shape differences in hemodynamic responses at the group level. Frontiers in Neuroscience 9. https://doi.org/10.3389/fnins.2015.00375

31. Chen, G., Saad, Z.S., Nath, A.R., Beauchamp, M.S., Cox, R.W., 2012. FMRI group analysis combining effect estimates and their variances. Neuroimage 60, 747–765. https://doi.org/10.1016/j.neuroimage.2011.12.060

32. Chen, J.E., Jahanian, H., Glover, G.H., 2016. Nuisance Regression of High-Frequency Functional Magnetic Resonance Imaging Data: Denoising Can Be Noisy. Brain Connectivity 7, 13–24. https://doi.org/10.1089/brain.2016.0441

33. Chen, J.E., Polimeni, J.R., Bollmann, S., Glover, G.H., 2019. On the analysis of rapidly sampled fMRI data. NeuroImage 188, 807–820. https://doi.org/10.1016/j.neuroimage.2019.02.008

34. Chen, L., T. Vu, A., Xu, J., Moeller, S., Ugurbil, K., Yacoub, E., Feinberg, D.A., 2015. Evaluation of highly accelerated simultaneous multi-slice EPI for fMRI. NeuroImage 104, 452–459. https://doi.org/10.1016/j.neuroimage.2014.10.027

35. Cichy, R.M., Pantazis, D., Oliva, A., 2016. Similarity-Based Fusion of MEG and fMRI Reveals Spatio-Temporal Dynamics in Human Cortex During Visual Object Recognition. Cerebral Cortex 26, 3563–3579. https://doi.org/10.1093/cercor/bhw135

36. Corbin, N., Todd, N., Friston, K.J., Callaghan, M.F., 2018. Accurate modeling of temporal correlations in rapidly sampled fMRI time series. Human Brain Mapping 39, 3884–3897. https://doi.org/10.1002/hbm.24218

37. Cox, R.W., Chen, G., Glen, D.R., Reynolds, R.C., Taylor, P.A., 2017. FMRI Clustering in AFNI: False-Positive Rates Redux. Brain Connect 7, 152–171. https://doi.org/10.1089/brain.2016.0475

38. Dale, A.M., 1999. Optimal experimental design for event-related fMRI. Human Brain Mapping 8, 109–114. https://doi.org/10.1002/(SICI)1097-0193(1999)8:2/3<109::AID-HBM7>3.0.CO;2-W

39. De Martino, F., Moerel, M., Ugurbil, K., Formisano, E., Yacoub, E., 2015. Less noise, more activation: Multiband acquisition schemes for auditory functional MRI. Magn Reson Med 74, 462–467. https://doi.org/10.1002/mrm.25408

40. De Martino, F., Yacoub, E., Kemper, V., Moerel, M., Uludağ, K., De Weerd, P., Ugurbil, K., Goebel, R., Formisano, E., 2018. The impact of ultra-high field MRI on cognitive and computational neuroimaging. NeuroImage, Neuroimaging with Ultra-high Field MRI: Present and Future 168, 366–382. https://doi.org/10.1016/j.neuroimage.2017.03.060

41. de Zwart, J.A., Silva, A.C., van Gelderen, P., Kellman, P., Fukunaga, M., Chu, R., Koretsky, A.P., Frank, J.A., Duyn, J.H., 2005. Temporal dynamics of the BOLD fMRI impulse response. NeuroImage 24, 667–677. https://doi.org/10.1016/j.neuroimage.2004.09.013

42. Debener, S., Mullinger, K.J., Niazy, R.K., Bowtell, R.W., 2008. Properties of the ballistocardiogram artefact as revealed by EEG recordings at 1.5, 3 and 7 T static magnetic field strength. International Journal of Psychophysiology, Integration of EEG and fMRI 67, 189–199. https://doi.org/10.1016/j.ijpsycho.2007.05.015

43. Deligianni, F., Centeno, M., Carmichael, D.W., Clayden, J.D., 2014. Relating resting-state fMRI and EEG whole-brain connectomes across frequency bands. Front. Neurosci. 0. https://doi.org/10.3389/fnins.2014.00258

44. Demetriou, L., Kowalczyk, O.S., Tyson, G., Bello, T., Newbould, R.D., Wall, M.B., 2018. A comprehensive evaluation of increasing temporal resolution with multiband-accelerated protocols and effects on statistical outcome measures in fMRI. NeuroImage 176, 404–416. https://doi.org/10.1016/j.neuroimage.2018.05.011

45. Dietrich, O., Raya, J.G., Reeder, S.B., Reiser, M.F., Schoenberg, S.O., 2007. Measurement of signal-to-noise ratios in MR images: Influence of multichannel coils, parallel imaging, and reconstruction filters. Journal of Magnetic Resonance Imaging 26, 375–385. https://doi.org/10.1002/jmri.20969

46. Dowdle, L.T., Ghose, G., Ugurbil, K., Yacoub, E., Vizioli, L., 2021. Clarifying the role of higher-level cortices in resolving perceptual ambiguity using ultra high field fMRI. NeuroImage 227, 117654. https://doi.org/10.1016/j.neuroimage.2020.117654

47. Drew, P.J., 2019. Vascular and neural basis of the BOLD signal. Curr Opin Neurobiol 58, 61–69. https://doi.org/10.1016/j.conb.2019.06.004

48. Durand, E., van de Moortele, P.-F., Pachot-Clouard, M., Bihan, D.L., 2001. Artifact due to B0 fluctuations in fMRI: Correction using the k-space central line. Magnetic Resonance in Medicine 46, 198–201. https://doi.org/10.1002/mrm.1177

49. Eklund, A., Nichols, T.E., Knutsson, H., 2016. Cluster failure: Why fMRI inferences for spatial extent have inflated false-positive rates. PNAS 113, 7900–7905. https://doi.org/10.1073/pnas.1602413113

50. Eklund Anders, E., Anderson, M., Josephson, M, J., H, K., 2012. Does parametric fMRI analysis with SPM yield valid results? An empirical study of 1484 rest datasets. Neuroimage 61, 565–578. https://doi.org/10.1016/j.neuroimage.2012.03.093

51. Erdoğan, S.B., Tong, Y., Hocke, L.M., Lindsey, K.P., deB Frederick, B., 2016. Correcting for Blood Arrival Time in Global Mean Regression Enhances Functional Connectivity Analysis of Resting State fMRI-BOLD Signals. Front. Hum. Neurosci. 10. https://doi.org/10.3389/fnhum.2016.00311

52. Fair, D.A., Miranda-Dominguez, O., Snyder, A.Z., Perrone, A., Earl, E.A., Van, A.N., Koller, J.M., Feczko, E., Tisdall, M.D., van der Kouwe, A., Klein, R.L., Mirro, A.E., Hampton, J.M., Adeyemo, B., Laumann, T.O., Gratton, C., Greene, D.J., Schlaggar, B.L., Hagler, D.J., Watts, R., Garavan, H., Barch, D.M., Nigg, J.T., Petersen, S.E., Dale, A.M., Feldstein-Ewing, S.W., Nagel, B.J., Dosenbach, N.U.F., 2020. Correction of respiratory artifacts in MRI head motion estimates. NeuroImage 208, 116400. https://doi.org/10.1016/j.neuroimage.2019.116400

53. Feinberg, D.A., Moeller, S., Smith, S.M., Auerbach, E., Ramanna, S., Glasser, M.F., Miller, K.L., Ugurbil, K., Yacoub, E., 2010. Multiplexed Echo Planar Imaging for Sub-Second Whole Brain FMRI and Fast Diffusion Imaging. PLOS ONE 5, e15710. https://doi.org/10.1371/journal.pone.0015710

54. Fischl, B., Dale, A.M., 2000. Measuring the thickness of the human cerebral cortex from magnetic resonance images. PNAS 97, 11050–11055. https://doi.org/10.1073/pnas.200033797

55. Formisano, E., Goebel, R., 2003. Tracking cognitive processes with functional MRI mental chronometry. Curr. Opin. Neurobiol. 13, 174–181.

56. Formisano, E., Linden, D.E.J., Di Salle, F., Trojano, L., Esposito, F., Sack, A.T., Grossi, D., Zanella, F.E., Goebel, R., 2002. Tracking the Mind’s Image in the Brain I: Time-Resolved fMRI during Visuospatial Mental Imagery. Neuron 35, 185–194. https://doi.org/10.1016/S0896-6273(02)00747-X

57. Fransson, P., Krüger, G., Merboldt, K.D., Frahm, J., 1999. Temporal and spatial MRI responses to subsecond visual activation. Magnetic Resonance Imaging 17, 1–7. https://doi.org/10.1016/S0730-725X(98)00163-5

58. Fransson, P., Krüger, G., Merboldt, K.D., Frahm, J., 1998. Temporal characteristics of oxygenation-sensitive MRI responses to visual activation in humans. Magn Reson Med 39, 912–919. https://doi.org/10.1002/mrm.1910390608

59. Friston, K.J., Fletcher, P., Josephs, O., Holmes, A., Rugg, M.D., Turner, R., 1998. Event-Related fMRI: Characterizing Differential Responses. NeuroImage 7, 30–40. https://doi.org/10.1006/nimg.1997.0306

60. Friston, K.J., Glaser, D.E., Henson, R.N.A., Kiebel, S., Phillips, C., Ashburner, J., 2002. Classical and Bayesian Inference in Neuroimaging: Applications. NeuroImage 16, 484–512. https://doi.org/10.1006/nimg.2002.1091

61. Geissler, A., Lanzenberger, R., Barth, M., Tahamtan, A.R., Milakara, D., Gartus, A., Beisteiner, R., 2005. Influence of fMRI smoothing procedures on replicability of fine scale motor localization. Neuroimage 24, 323–331. https://doi.org/10.1016/j.neuroimage.2004.08.042

62. Gentile, F., Ales, J., Rossion, B., 2017. Being BOLD: The neural dynamics of face perception. Human Brain Mapping 38, 120–139. https://doi.org/10.1002/hbm.23348

63. Glasser, M.F., Smith, S.M., Marcus, D.S., Andersson, J., Auerbach, E.J., Behrens, T.E.J., Coalson, T.S., Harms, M.P., Jenkinson, M., Moeller, S., Robinson, E.C., Sotiropoulos, S.N., Xu, J., Yacoub, E., Ugurbil, K., Van Essen, D.C., 2016. The Human Connectome Project’s Neuroimaging Approach. Nat Neurosci 19, 1175– 1187. https://doi.org/10.1038/nn.4361

64. Glover, G.H., 1999. Deconvolution of Impulse Response in Event-Related BOLD fMRI1. NeuroImage 9, 416– 429. https://doi.org/10.1006/nimg.1998.0419

65. Glover, G.H., Li, T.-Q., Ress, D., 2000. Image-based method for retrospective correction of physiological motion effects in fMRI: RETROICOR. Magnetic Resonance in Medicine 44, 162–167. https://doi.org/10.1002/1522-2594(200007)44:1<162::AID-MRM23>3.0.CO;2-E

66. Gohel, S.R., Biswal, B.B., 2014. Functional Integration Between Brain Regions at Rest Occurs in Multiple-Frequency Bands. Brain Connectivity 5, 23–34. https://doi.org/10.1089/brain.2013.0210

67. Goldman, R.I., Stern, J.M., Engel, J.J., Cohen, M.S., 2002. Simultaneous EEG and fMRI of the alpha rhythm. NeuroReport 13, 2487–2492.

68. Gonzalez-Castillo, J., Saad, Z.S., Handwerker, D.A., Inati, S.J., Brenowitz, N., Bandettini, P.A., 2012. Whole-brain, time-locked activation with simple tasks revealed using massive averaging and model-free analysis. Proceedings of the National Academy of Sciences of the United States of America 109, 5487– 5492. https://doi.org/10.1073/pnas.1121049109

69. Goodyear, B.G., Menon, R.S., 2001. Brief visual stimulation allows mapping of ocular dominance in visual cortex using fMRI. Hum Brain Mapp 14, 210–217. https://doi.org/10.1002/hbm.1053

70. Griffanti, L., Salimi-Khorshidi, G., Beckmann, C.F., Auerbach, E.J., Douaud, G., Sexton, C.E., Zsoldos, E., Ebmeier, K.P., Filippini, N., Mackay, C.E., Moeller, S., Xu, J., Yacoub, E., Baselli, G., Ugurbil, K., Miller, K.L., Smith, S.M., 2014. ICA-based artefact removal and accelerated fMRI acquisition for improved resting state network imaging. NeuroImage 95, 232–247. https://doi.org/10.1016/j.neuroimage.2014.03.034

71. Handwerker, D.A., Ollinger, J.M., D’Esposito, M., 2004. Variation of BOLD hemodynamic responses across subjects and brain regions and their effects on statistical analyses. NeuroImage 21, 1639–1651. https://doi.org/10.1016/j.neuroimage.2003.11.029

72. Haxby, J.V., Connolly, A.C., Guntupalli, J.S., 2014. Decoding Neural Representational Spaces Using Multivariate Pattern Analysis. Annual Review of Neuroscience 37, 435–456. https://doi.org/10.1146/annurev-neuro-062012-170325

73. Haxby, J.V., Gobbini, M.I., Furey, M.L., Ishai, A., Schouten, J.L., Pietrini, P., 2001. Distributed and overlapping representations of faces and objects in ventral temporal cortex. Science 293, 2425–2430. https://doi.org/10.1126/science.1063736

74. Hayasaka, S., Nichols, T.E., 2003. Validating cluster size inference: random field and permutation methods. Neuroimage 20, 2343–2356. https://doi.org/10.1016/j.neuroimage.2003.08.003

75. Henson, R., Rugg, M.D., Friston, K.J., 2001. The choice of basis functions in event-related fMRI. NeuroImage, Originally published as Volume 13, Number 6, Part 2 13, 149. https://doi.org/10.1016/S1053-8119(01)91492-2

76. Henson, R.N.A., Price, C.J., Rugg, M.D., Turner, R., Friston, K.J., 2002. Detecting Latency Differences in Event-Related BOLD Responses: Application to Words versus Nonwords and Initial versus Repeated Face Presentations. NeuroImage 15, 83–97. https://doi.org/10.1006/nimg.2001.0940

77. Hernandez, L., Badre, D., Noll, D., Jonides, J., 2002. Temporal Sensitivity of Event-Related fMRI. NeuroImage 17, 1018–1026. https://doi.org/10.1006/nimg.2001.1017

78. Hill, P.F., Seger, S.E., Yoo, H.B., King, D.R., Wang, D.X., Lega, B.C., Rugg, M.D., 2021. Distinct Neurophysiological Correlates of the fMRI BOLD Signal in the Hippocampus and Neocortex. J. Neurosci. 41, 6343–6352. https://doi.org/10.1523/JNEUROSCI.0278-21.2021

79. Hocke, L.M., Tong, Y., Lindsey, K.P., de Frederick, B. B., 2016. Comparison of peripheral near-infrared spectroscopy low-frequency oscillations to other denoising methods in resting state functional MRI with ultrahigh temporal resolution. Magnetic Resonance in Medicine 76, 1697–1707. https://doi.org/10.1002/mrm.26038

80. Hu, X., Le, T.H., U urbil, K., 1997. Evaluation of the early response in fMRI in individual subjects using short stimulus duration. Magnetic Resonance in Medicine 37, 877–884. https://doi.org/10.1002/mrm.1910370612

81. Hu, X., Yacoub, E., 2012. The story of the initial dip in fMRI. Neuroimage 62, 1103–1108. https://doi.org/10.1016/j.neuroimage.2012.03.005

82. Hutchison, R.M., Womelsdorf, T., Allen, E.A., Bandettini, P.A., Calhoun, V.D., Corbetta, M., Della Penna, S., Duyn, J.H., Glover, G.H., Gonzalez-Castillo, J., Handwerker, D.A., Keilholz, S., Kiviniemi, V., Leopold, D.A., de Pasquale, F., Sporns, O., Walter, M., Chang, C., 2013. Dynamic functional connectivity: Promise, issues, and interpretations. NeuroImage, Mapping the Connectome 80, 360–378. https://doi.org/10.1016/j.neuroimage.2013.05.079

83. Jacobs, J., Stich, J., Zahneisen, B., Assländer, J., Ramantani, G., Schulze-Bonhage, A., Korinthenberg, R., Hennig, J., LeVan, P., 2014. Fast fMRI provides high statistical power in the analysis of epileptic networks. Neuroimage 88, 282–294. https://doi.org/10.1016/j.neuroimage.2013.10.018

84. Jäger, V., Dümpelmann, M., LeVan, P., Ramantani, G., Mader, I., Schulze-Bonhage, A., Jacobs, J., 2015. Concordance of Epileptic Networks Associated with Epileptic Spikes Measured by High-Density EEG and Fast fMRI. PLOS ONE 10, e0140537. https://doi.org/10.1371/journal.pone.0140537

85. Jo, H.J., Lee, J.-M., Kim, J.-H., Choi, C.-H., Gu, B.-M., Kang, D.-H., Ku, J., Kwon, J.S., Kim, S.I., 2008. Artificial shifting of fMRI activation localized by volume- and surface-based analyses. NeuroImage 40, 1077– 1089. https://doi.org/10.1016/j.neuroimage.2007.12.036

86. Jorge, J., Grouiller, F., Ipek, Ö., Stoermer, R., Michel, C.M., Figueiredo, P., van der Zwaag, W., Gruetter, R., 2015. Simultaneous EEG–fMRI at ultra-high field: Artifact prevention and safety assessment. NeuroImage 105, 132–144. https://doi.org/10.1016/j.neuroimage.2014.10.055

87. Kaneoke, Y., Donishi, T., Iwatani, J., Ukai, S., Shinosaki, K., Terada, M., 2012. Variance and Autocorrelation of the Spontaneous Slow Brain Activity. PLOS ONE 7, e38131. https://doi.org/10.1371/journal.pone.0038131

88. Kasper, L., Bollmann, S., Diaconescu, A.O., Hutton, C., Heinzle, J., Iglesias, S., Hauser, T.U., Sebold, M., Manjaly, Z.-M., Pruessmann, K.P., Stephan, K.E., 2017. The PhysIO Toolbox for Modeling Physiological Noise in fMRI Data. Journal of Neuroscience Methods 276, 56–72. https://doi.org/10.1016/j.jneumeth.2016.10.019

89. Kay, K., Jamison, K.W., Zhang, R.-Y., Uğurbil, K., 2020. A temporal decomposition method for identifying venous effects in task-based fMRI. Nature Methods 17, 1033–1039. https://doi.org/10.1038/s41592-020-0941-6

90. Kay, K., Rokem, A., Winawer, J., Dougherty, R., Wandell, B., 2013. GLMdenoise: a fast, automated technique for denoising task-based fMRI data. Front. Neurosci. 7. https://doi.org/10.3389/fnins.2013.00247

91. Kay, K.N., David, S.V., Prenger, R.J., Hansen, K.A., Gallant, J.L., 2008. Modeling low-frequency fluctuation and hemodynamic response timecourse in event-related fMRI. Human Brain Mapping 29, 142–156. https://doi.org/10.1002/hbm.20379

92. Kim, S.-G., Richter, W., Uurbil, K., 1997. Limitations of temporal resolution in functional MRI. Magnetic Resonance in Medicine 37, 631–636. https://doi.org/10.1002/mrm.1910370427

93. Kirilina, E., Lutti, A., Poser, B.A., Blankenburg, F., Weiskopf, N., 2016. The quest for the best: The impact of different EPI sequences on the sensitivity of random effect fMRI group analyses. Neuroimage 126, 49– 59. https://doi.org/10.1016/j.neuroimage.2015.10.071

94. Koopmans, P.J., Barth, M., Norris, D.G., 2010. Layer-specific BOLD activation in human V1. Hum Brain Mapp 31, 1297–1304. https://doi.org/10.1002/hbm.20936

95. Koopmans, P.J., Barth, M., Orzada, S., Norris, D.G., 2011. Multi-echo fMRI of the cortical laminae in humans at 7T. NeuroImage 56, 1276–1285. https://doi.org/10.1016/j.neuroimage.2011.02.042

96. Koopmans, P.J., Pfaffenrot, V., 2021. Enhanced POCS reconstruction for partial Fourier imaging in multi-echo and time-series acquisitions. Magnetic Resonance in Medicine 85, 140–151. https://doi.org/10.1002/mrm.28417

97. Kriegeskorte, N., Bandettini, P., 2007. Analyzing for information, not activation, to exploit high-resolution fMRI. NeuroImage 38, 649–662. https://doi.org/10.1016/j.neuroimage.2007.02.022

98. Kriegeskorte, N., Goebel, R., Bandettini, P., 2006. Information-based functional brain mapping. PNAS 103, 3863–3868. https://doi.org/10.1073/pnas.0600244103

99. Kundu, P., Inati, S.J., Evans, J.W., Luh, W.M., Bandettini, P.A., 2012. Differentiating BOLD and non-BOLD signals in fMRI time series using multi-echo EPI. Neuroimage 60, 1759–70. https://doi.org/10.1016/j.neuroimage.2011.12.028

100. Kundu, P., Voon, V., Balchandani, P., Lombardo, M.V., Poser, B.A., Bandettini, P., 2017. Multi-Echo fMRI: A Review of Applications in fMRI Denoising and Analysis of BOLD Signals. Neuroimage. https://doi.org/10.1016/j.neuroimage.2017.03.033

101. Larkman, D.J., Hajnal, J.V., Herlihy, A.H., Coutts, G.A., Young, I.R., Ehnholm, G., 2001. Use of multicoil arrays for separation of signal from multiple slices simultaneously excited. J Magn Reson Imaging 13, 313–317. https://doi.org/10.1002/1522-2586(200102)13:2<313::aid-jmri1045>3.0.co;2-w

102. Le, T.H., Hu, X., 1996. Retrospective estimation and correction of physiological artifacts in fMRI by direct extraction of physiological activity from MR data. Magn Reson Med 35, 290–298. https://doi.org/10.1002/mrm.1910350305

103. Lenoski, B., Baxter, L.C., Karam, L.J., Maisog, J., Debbins, J., 2008. On the Performance of Autocorrelation Estimation Algorithms for fMRI Analysis. IEEE Journal of Selected Topics in Signal Processing 2, 828– 838. https://doi.org/10.1109/JSTSP.2008.2007819

104. Lewis, L.D., Setsompop, K., Rosen, B.R., Polimeni, J.R., 2018. Stimulus-dependent hemodynamic response timing across the human subcortical-cortical visual pathway identified through high spatiotemporal resolution 7T fMRI. NeuroImage 181, 279–291. https://doi.org/10.1016/j.neuroimage.2018.06.056

105. Lewis, L.D., Setsompop, K., Rosen, B.R., Polimeni, J.R., 2016. Fast fMRI can detect oscillatory neural activity in humans. Proceedings of the National Academy of Sciences 113, E6679–E6685. https://doi.org/10.1073/pnas.1608117113

106. Liao, C.H., Worsley, K.J., Poline, J.-B., Aston, J.A.D., Duncan, G.H., Evans, A.C., 2002. Estimating the Delay of the fMRI Response. NeuroImage 16, 593–606. https://doi.org/10.1006/nimg.2002.1096

107. Lin, F.H., Polimeni, J.R., Lin, J.F.L., Tsai, K.W.K., Chu, Y.H., Wu, P.Y., Li, Y.T., Hsu, Y.C., Tsai, S.Y., Kuo, W.J., 2018. Relative latency and temporal variability of hemodynamic responses at the human primary visual cortex. NeuroImage, Pushing the spatio-temporal limits of MRI and fMRI 164, 194–201. https://doi.org/10.1016/j.neuroimage.2017.01.041

108. Lin, F.-H., Polimeni, J.R., Tsai, K.W.-K., Witzel, T., Chang, W.-T., Kuo, W.-J., Belliveau, J.W., 2011. The limit of relative timing accuracy of BOLD fMRI in human visual cortex 1.

109. Lin, F.H., Witzel, T., Raij, T., Ahveninen, J., Wen-Kai Tsai, K., Chu, Y.H., Chang, W.T., Nummenmaa, A., Polimeni, J.R., Kuo, W.J., Hsieh, J.C., Rosen, B.R., Belliveau, J.W., 2013. FMRI hemodynamics accurately reflects neuronal timing in the human brain measured by MEG. NeuroImage 78, 372–384. https://doi.org/10.1016/j.neuroimage.2013.04.017

110. Lindquist, M.A., Geuter, S., Wager, T.D., Caffo, B.S., 2019. Modular preprocessing pipelines can reintroduce artifacts into fMRI data. Human Brain Mapping 40, 2358–2376. https://doi.org/10.1002/hbm.24528

111. Lindquist, M.A., Loh, J.M., Atlas, L.Y., Wager, T.D., 2009. Modeling the Hemodynamic Response Function in fMRI: Efficiency, Bias and Mis-modeling. Neuroimage 45, S187–S198. https://doi.org/10.1016/j.neuroimage.2008.10.065

112. Logothetis, N.K., Pauls, J., Augath, M., Trinath, T., Oeltermann, A., 2001. Neurophysiological investigation of the basis of the fMRI signal. Nature 412, 150–157. https://doi.org/10.1038/35084005

113. Luo, Q., Misaki, M., Mulyana, B., Wong, C.-K., Bodurka, J., 2020. Improved autoregressive model for correction of noise serial correlation in fast fMRI. Magnetic Resonance in Medicine 84, 1293–1305. https://doi.org/10.1002/mrm.28203

114. Margalit, E., Jamison, K.W., Weiner, K.S., Vizioli, L., Zhang, R.-Y., Kay, K.N., Grill-Spector, K., 2020. Ultra-high-resolution fMRI of human ventral temporal cortex reveals differential representation of categories and domains. J. Neurosci. https://doi.org/10.1523/JNEUROSCI.2106-19.2020

115. Masterton, R.A.J., Abbott, D.F., Fleming, S.W., Jackson, G.D., 2007. Measurement and reduction of motion and ballistocardiogram artefacts from simultaneous EEG and fMRI recordings. NeuroImage 37, 202–211. https://doi.org/10.1016/j.neuroimage.2007.02.060

116. Mayhew, S.D., Bagshaw, A.P., 2017. Dynamic spatiotemporal variability of alpha-BOLD relationships during the resting-state and task-evoked responses. NeuroImage 155, 120–137. https://doi.org/10.1016/j.neuroimage.2017.04.051

117. McDowell, A.R., Carmichael, D.W., 2019. Optimal repetition time reduction for single subject event-related functional magnetic resonance imaging. Magnetic Resonance in Medicine 81, 1890–1897. https://doi.org/10.1002/mrm.27498

118. McGuire, J.T., Kable, J.W., 2015. Medial prefrontal cortical activity reflects dynamic re-evaluation during voluntary persistence. Nature Neuroscience 18, 760–766. https://doi.org/10.1038/nn.3994

119. Menon, R.S., Kim, S.-G., 1999. Spatial and temporal limits in cognitive neuroimaging with fMRI. Trends in Cognitive Sciences 3, 207–216. https://doi.org/10.1016/S1364-6613(99)01329-7

120. Menon, R.S., Luknowsky, D.C., Gati, J.S., 1998. Mental chronometry using latency-resolved functional MRI. PNAS 95, 10902–10907.

121. Menon, R.S., Ogawa, S., Hu, X., Strupp, J.P., Anderson, P., U urbil, K., 1995. BOLD Based Functional MRI at 4 Tesla Includes a Capillary Bed Contribution: Echo-Planar Imaging Correlates with Previous Optical Imaging Using Intrinsic Signals. Magnetic Resonance in Medicine 33, 453–459. https://doi.org/10.1002/mrm.1910330323

122. Meyer, M.C., Janssen, R.J., van Oort, E.S.B., Beckmann, C.F., Barth, M., 2013. The Quest for EEG Power Band Correlation with ICA Derived fMRI Resting State Networks. Front. Hum. Neurosci. 0. https://doi.org/10.3389/fnhum.2013.00315

123. Miezin, F.M., Maccotta, L., Ollinger, J.M., Petersen, S.E., Buckner, R.L., 2000. Characterizing the Hemodynamic Response: Effects of Presentation Rate, Sampling Procedure, and the Possibility of Ordering Brain Activity Based on Relative Timing. NeuroImage 11, 735–759. https://doi.org/10.1006/nimg.2000.0568

124. Moeller, S., Yacoub, E., Olman, C.A., Auerbach, E., Strupp, J., Harel, N., Uğurbil, K., 2010. Multiband multislice GE-EPI at 7 tesla, with 16-fold acceleration using partial parallel imaging with application to high spatial and temporal whole-brain fMRI. Magnetic Resonance in Medicine 63, 1144–1153. https://doi.org/10.1002/mrm.22361

125. Moia, S., Termenon, M., Uruñuela, E., Chen, G., Stickland, R.C., Bright, M.G., Caballero-Gaudes, C., 2021. ICA-based denoising strategies in breath-hold induced cerebrovascular reactivity mapping with multi echo BOLD fMRI. NeuroImage 233, 117914. https://doi.org/10.1016/j.neuroimage.2021.117914

126. Morgan, A.T., Nothnagel, N., Petro, L.S., Goense, J., Muckli, L., 2020. High-resolution line-scanning reveals distinct visual response properties across human cortical layers. bioRxiv 2020.06.30.179762. https://doi.org/10.1101/2020.06.30.179762

127. Murta, T., Leite, M., Carmichael, D.W., Figueiredo, P., Lemieux, L., 2015. Electrophysiological correlates of the BOLD signal for EEG-informed fMRI. Human Brain Mapping 36, 391–414. https://doi.org/10.1002/hbm.22623

128. Neuner, I., Arrubla, J., Felder, J., Shah, N.J., 2014. Simultaneous EEG–fMRI acquisition at low, high and ultra-high magnetic fields up to 9.4T: Perspectives and challenges. NeuroImage, Multimodal Data Fusion 102, 71–79. https://doi.org/10.1016/j.neuroimage.2013.06.048

129. Nichols, T.E., 2012. Multiple testing corrections, nonparametric methods, and random field theory. NeuroImage, 20 YEARS OF fMRI 62, 811–815. https://doi.org/10.1016/j.neuroimage.2012.04.014

130. Ogawa, S., Lee, T.M., Kay, A.R., Tank, D.W., 1990. Brain magnetic resonance imaging with contrast dependent on blood oxygenation. Proc Natl Acad Sci U S A 87, 9868–9872.

131. Ogawa, S., Lee, T.-M., Stepnoski, R., Chen, W., Zhu, X.-H., Ugurbil, K., 2000. An approach to probe some neural systems interaction by functional MRI at neural time scale down to milliseconds. PNAS 97, 11026– 11031. https://doi.org/10.1073/pnas.97.20.11026

132. Olafsson, V., Kundu, P., Wong, E.C., Bandettini, P.A., Liu, T.T., 2015. Enhanced identification of BOLD-like components with multi-echo simultaneous multi-slice (MESMS) fMRI and multi-echo ICA. Neuroimage 112, 43–51. https://doi.org/10.1016/j.neuroimage.2015.02.052

133. Ollinger, J.M., Shulman, G.L., Corbetta, M., 2001. Separating Processes within a Trial in Event-Related Functional MRI: I. The Method. NeuroImage 13, 210–217. https://doi.org/10.1006/nimg.2000.0710

134. Olman, C.A., Harel, N., Feinberg, D.A., He, S., Zhang, P., Ugurbil, K., Yacoub, E., 2012. Layer-Specific fMRI Reflects Different Neuronal Computations at Different Depths in Human V1. PLoS One 7. https://doi.org/10.1371/journal.pone.0032536

135. Olszowy, W., Aston, J., Rua, C., Williams, G.B., 2019. Accurate autocorrelation modeling substantially improves fMRI reliability. Nature Communications 10, 1–11. https://doi.org/10.1038/s41467-019-09230-w

136. Park, B., Shim, W.M., James, O., Park, H., 2019. Possible links between the lag structure in visual cortex and visual streams using fMRI. Sci Rep 9, 4283. https://doi.org/10.1038/s41598-019-40728-x

137. Pernet, C.R., 2014. Misconceptions in the use of the General Linear Model applied to functional MRI: a tutorial for junior neuro-imagers. Front Neurosci 8. https://doi.org/10.3389/fnins.2014.00001

138. Petridou, N., Siero, J.C.W., 2019. Laminar fMRI: What can the time domain tell us? NeuroImage 197, 761–771. https://doi.org/10.1016/j.neuroimage.2017.07.040

139. Polimeni, J.R., Lewis, L.D., 2021. Imaging faster neural dynamics with fast fMRI: a need for updated models of the hemodynamic response. Progress in Neurobiology.

140. Posner, M.I., 1978. Chronometric explorations of mind, Chronometric explorations of mind. Lawrence Erlbaum, Oxford, England.

141. Posse, S., Ackley, E., Mutihac, R., Rick, J., Shane, M., Murray-Krezan, C., Zaitsev, M., Speck, O., 2012. Enhancement of temporal resolution and BOLD sensitivity in real-time fMRI using multi-slab echo-volumar imaging. Neuroimage 61, 115–130. https://doi.org/10.1016/j.neuroimage.2012.02.059

142. Power, J.D., Lynch, C.J., Silver, B.M., Dubin, M.J., Martin, A., Jones, R.M., 2019. Distinctions among real and apparent respiratory motions in human fMRI data. NeuroImage 201, 116041. https://doi.org/10.1016/j.neuroimage.2019.116041

143. Power, J.D., Plitt, M., Kundu, P., Bandettini, P.A., Martin, A., 2017. Temporal interpolation alters motion in fMRI scans: Magnitudes and consequences for artifact detection. PLoS One 12, e0182939. https://doi.org/10.1371/journal.pone.0182939

144. Preti, M.G., Bolton, T.A., Van De Ville, D., 2017. The dynamic functional connectome: State-of-the-art and perspectives. NeuroImage, Functional Architecture of the Brain 160, 41–54. https://doi.org/10.1016/j.neuroimage.2016.12.061

145. Puckett, A.M., Aquino, K.M., Robinson, P.A., Breakspear, M., Schira, M.M., 2016. The spatiotemporal hemodynamic response function for depth-dependent functional imaging of human cortex. NeuroImage 139, 240–248. https://doi.org/10.1016/j.neuroimage.2016.06.019

146. Purdon, P.L., Weisskoff, R.M., 1998. Effect of temporal autocorrelation due to physiological noise and stimulus paradigm on voxel-level false-positive rates in fMRI. Hum Brain Mapp 6, 239–249.

147. Raj, D., Anderson, A.W., Gore, J.C., 2001. Respiratory effects in human functional magnetic resonance imaging due to bulk susceptibility changes. Phys. Med. Biol. 46, 3331–3340. https://doi.org/10.1088/0031-9155/46/12/318

148. Ramon, M., Vizioli, L., Liu-Shuang, J., Rossion, B., 2015. Neural microgenesis of personally familiar face recognition. PNAS 112, E4835–E4844. https://doi.org/10.1073/pnas.1414929112

149. Rosa, M.J., Kilner, J., Blankenburg, F., Josephs, O., Penny, W., 2010. Estimating the transfer function from neuronal activity to BOLD using simultaneous EEG-fMRI. Neuroimage 49, 1496–1509. https://doi.org/10.1016/j.neuroimage.2009.09.011

150. Sahib, A.K., Erb, M., Marquetand, J., Martin, P., Elshahabi, A., Klamer, S., Vulliemoz, S., Scheffler, K., Ethofer, T., Focke, N.K., 2018. Evaluating the impact of fast-fMRI on dynamic functional connectivity in an event-based paradigm. PLOS ONE 13, e0190480. https://doi.org/10.1371/journal.pone.0190480

151. Sahib, A.K., Mathiak, K., Erb, M., Elshahabi, A., Klamer, S., Scheffler, K., Focke, N.K., Ethofer, T., 2016. Effect of temporal resolution and serial autocorrelations in event-related functional MRI. Magnetic Resonance in Medicine 76, 1805–1813. https://doi.org/10.1002/mrm.26073

152. Salek-Haddadi, A., Diehl, B., Hamandi, K., Merschhemke, M., Liston, A., Friston, K., Duncan, J.S., Fish, D.R., Lemieux, L., 2006. Hemodynamic correlates of epileptiform discharges: An EEG-fMRI study of 63 patients with focal epilepsy. Brain Research 1088, 148–166. https://doi.org/10.1016/j.brainres.2006.02.098

153. Shmuel, A., Yacoub, E., Chaimow, D., Logothetis, N.K., Ugurbil, K., 2007. Spatio-temporal point-spread function of fMRI signal in human gray matter at 7 Tesla. NeuroImage 35, 539–552. https://doi.org/10.1016/j.neuroimage.2006.12.030

154. Siero, J.C., Petridou, N., Hoogduin, H., Luijten, P.R., Ramsey, N.F., 2011. Cortical Depth-Dependent Temporal Dynamics of the BOLD Response in the Human Brain. J Cereb Blood Flow Metab 31, 1999–2008. https://doi.org/10.1038/jcbfm.2011.57

155. Siero, J.C.W., Hendrikse, J., Hoogduin, H., Petridou, N., Luijten, P., Donahue, M.J., 2015. Cortical depth dependence of the BOLD initial dip and poststimulus undershoot in human visual cortex at 7 Tesla. Magnetic Resonance in Medicine 73, 2283–2295. https://doi.org/10.1002/mrm.25349

156. Siero, J.C.W., Ramsey, N.F., Hoogduin, H., Klomp, D.W.J., Luijten, P.R., Petridou, N., 2013. BOLD Specificity and Dynamics Evaluated in Humans at 7 T: Comparing Gradient-Echo and Spin-Echo Hemodynamic Responses. PLOS ONE 8, e54560. https://doi.org/10.1371/journal.pone.0054560

157. Sigman, M., Jobert, A., LeBihan, D., Dehaene, S., 2007. Parsing a sequence of brain activations at psychological times using fMRI. NeuroImage 35, 655–668. https://doi.org/10.1016/j.neuroimage.2006.05.064

158. Silva, A.C., Koretsky, A.P., 2002. Laminar specificity of functional MRI onset times during somatosensory stimulation in rat. Proc Natl Acad Sci U S A 99, 15182–15187. https://doi.org/10.1073/pnas.222561899

159. Simony, E., Honey, C.J., Chen, J., Lositsky, O., Yeshurun, Y., Wiesel, A., Hasson, U., 2016. Dynamic reconfiguration of the default mode network during narrative comprehension. Nature Communications 7, 12141. https://doi.org/10.1038/ncomms12141

160. Sladky, R., Friston, K.J., Tröstl, J., Cunnington, R., Moser, E., Windischberger, C., 2011. Slice-timing effects and their correction in functional MRI. Neuroimage 58, 588–594. https://doi.org/10.1016/j.neuroimage.2011.06.078

161. Smith-Collins, A.P.R., Luyt, K., Heep, A., Kauppinen, R.A., 2015. High frequency functional brain networks in neonates revealed by rapid acquisition resting state fMRI. Human Brain Mapping 36, 2483–2494. https://doi.org/10.1002/hbm.22786

162. Stockmann, J.P., Wald, L.L., 2018. In vivo B0 field shimming methods for MRI at 7T. NeuroImage, Neuroimaging with Ultra-high Field MRI: Present and Future 168, 71–87. https://doi.org/10.1016/j.neuroimage.2017.06.013

163. T. Vu, A., Jamison, K., Glasser, M.F., Smith, S.M., Coalson, T., Moeller, S., Auerbach, E.J., Uğurbil, K., Yacoub, E., 2017. Tradeoffs in pushing the spatial resolution of fMRI for the 7T Human Connectome Project. NeuroImage, Cleaning up the fMRI time series: Mitigating noise with advanced acquisition and correction strategies 154, 23–32. https://doi.org/10.1016/j.neuroimage.2016.11.049

164. Taylor, A.J., Kim, J.H., Ress, D., 2018. Characterization of the hemodynamic response function across the majority of human cerebral cortex. NeuroImage 173, 322–331. https://doi.org/10.1016/j.neuroimage.2018.02.061

165. Thompson, S.K., Engel, S.A., Olman, C.A., 2014. Larger neural responses produce BOLD signals that begin earlier in time. Frontiers in Neuroscience 8. https://doi.org/10.3389/fnins.2014.00159

166. Todd, N., Josephs, O., Zeidman, P., Flandin, G., Moeller, S., Weiskopf, N., 2017. Functional Sensitivity of 2D Simultaneous Multi-Slice Echo-Planar Imaging: Effects of Acceleration on g-factor and Physiological Noise. Front. Neurosci. 11. https://doi.org/10.3389/fnins.2017.00158

167. Tong, Y., Hocke, L.M., Frederick, B. deB., 2014. Short Repetition Time Multiband Echo-Planar Imaging with Simultaneous Pulse Recording Allows Dynamic Imaging of the Cardiac Pulsation Signal. Magn Reson Med 72, 1268–1276. https://doi.org/10.1002/mrm.25041

168. Triantafyllou, C., Hoge, R.D., Krueger, G., Wiggins, C.J., Potthast, A., Wiggins, G.C., Wald, L.L., 2005. Comparison of physiological noise at 1.5 T, 3 T and 7 T and optimization of fMRI acquisition parameters. Neuroimage 26, 243–250. https://doi.org/10.1016/j.neuroimage.2005.01.007

169. Triantafyllou, C., Polimeni, J.R., Wald, L.L., 2011. Physiological noise and signal-to-noise ratio in fMRI with multi-channel array coils. Neuroimage 55, 597–606. https://doi.org/10.1016/j.neuroimage.2010.11.084

170. Uğurbil, K., Auerbach, E., Moeller, S., Grant, A., Wu, X., de Moortele, P.-F.V., Olman, C., DelaBarre, L., Schillak, S., Radder, J., Lagore, R., Adriany, G., 2019. Brain imaging with improved acceleration and SNR at 7 Tesla obtained with 64-channel receive array. Magnetic Resonance in Medicine 82, 495–509. https://doi.org/10.1002/mrm.27695

171. Uğurbil, K., Xu, J., Auerbach, E.J., Moeller, S., Vu, A.T., Duarte-Carvajalino, J.M., Lenglet, C., Wu, X., Schmitter, S., Van de Moortele, P.F., Strupp, J., Sapiro, G., De Martino, F., Wang, D., Harel, N., Garwood, M., Chen, L., Feinberg, D.A., Smith, S.M., Miller, K.L., Sotiropoulos, S.N., Jbabdi, S., Andersson, J.L.R., Behrens, T.E.J., Glasser, M.F., Van Essen, D.C., Yacoub, E., 2013. Pushing spatial and temporal resolution for functional and diffusion MRI in the Human Connectome Project. NeuroImage, Mapping the Connectome 80, 80–104. https://doi.org/10.1016/j.neuroimage.2013.05.012

172. van Gelderen, P., de Zwart, J.A., Starewicz, P., Hinks, R.S., Duyn, J.H., 2007. Real-time shimming to compensate for respiration-induced B0 fluctuations. Magn Reson Med 57, 362–368. https://doi.org/10.1002/mrm.21136

173. Vizioli, L., Bratch, A., Lao, J., Ugurbil, K., Muckli, L., Yacoub, E., 2018. Temporal multivariate pattern analysis (tMVPA): A single trial approach exploring the temporal dynamics of the BOLD signal. Journal of Neuroscience Methods 308, 74–87. https://doi.org/10.1016/j.jneumeth.2018.06.029

174. Vizioli, L., De Martino, F., Petro, L.S., Kersten, D., Ugurbil, K., Yacoub, E., Muckli, L., 2020a. Multivoxel Pattern of Blood Oxygen Level Dependent Activity can be sensitive to stimulus specific fine scale responses. Scientific Reports 10, 7565. https://doi.org/10.1038/s41598-020-64044-x

175. Vizioli, L., Moeller, S., Dowdle, L.T., Akçakaya, M., de Martino, F., Essa Yacoub, Ugurbil, K., 2020b. A Paradigm Change in Functional Brain Mapping: Suppressing the Thermal Noise in fMRI. bioRxiv 2020.11.04.368357. https://doi.org/10.1101/2020.11.04.368357

176. Vu, A.T., Beckett, A., Setsompop, K., Feinberg, D.A., 2018. Evaluation of SLIce Dithered Enhanced Resolution Simultaneous MultiSlice (SLIDER-SMS) for human fMRI. NeuroImage, Pushing the spatio-temporal limits of MRI and fMRI 164, 164–171. https://doi.org/10.1016/j.neuroimage.2017.02.001

177. Vu, A.T., Phillips, J.S., Kay, K., Phillips, M.E., Johnson, M.R., Shinkareva, S.V., Tubridy, S., Millin, R., Grossman, M., Gureckis, T., Bhattacharyya, R., Yacoub, E., 2016. Using precise word timing information improves decoding accuracy in a multiband-accelerated multimodal reading experiment. Cognitive Neuropsychology 33, 265–275. https://doi.org/10.1080/02643294.2016.1195343

178. Watanabe, M., Bartels, A., Macke, J.H., Murayama, Y., Logothetis, N.K., 2013. Temporal jitter of the BOLD signal reveals a reliable initial dip and improved spatial resolution. Curr. Biol. 23, 2146–2150. https://doi.org/10.1016/j.cub.2013.08.057

179. Weilke, F., Spiegel, S., Boecker, H., von Einsiedel, H.G., Conrad, B., Schwaiger, M., Erhard, P., 2001. Time-Resolved fMRI of Activation Patterns in M1 and SMA During Complex Voluntary Movement. Journal of Neurophysiology 85, 1858–1863. https://doi.org/10.1152/jn.2001.85.5.1858

180. White, T., O’Leary, D., Magnotta, V., Arndt, S., Flaum, M., Andreasen, N.C., 2001. Anatomic and functional variability: the effects of filter size in group fMRI data analysis. Neuroimage 13, 577–588. https://doi.org/10.1006/nimg.2000.0716

181. Whittingstall, K., Bartels, A., Singh, V., Kwon, S., Logothetis, N.K., 2010. Integration of EEG source imaging and fMRI during continuous viewing of natural movies. Magnetic Resonance Imaging, Proceedings of the International School on Magnetic Resonance and Brain Function 28, 1135–1142. https://doi.org/10.1016/j.mri.2010.03.042

182. Wirsich, J., Giraud, A.-L., Sadaghiani, S., 2020. Concurrent EEG- and fMRI-derived functional connectomes exhibit linked dynamics. NeuroImage 219, 116998. https://doi.org/10.1016/j.neuroimage.2020.116998

183. Worsley, K.J., Liao, C.H., Aston, J., Petre, V., Duncan, G.H., Morales, F., Evans, A.C., 2002. A general statistical analysis for fMRI data. Neuroimage 15, 1–15. https://doi.org/10.1006/nimg.2001.0933

184. Wu, X., Auerbach, E.J., Vu, A.T., Moeller, S., Van de Moortele, P.-F., Yacoub, E., Uğurbil, K., 2019. Human Connectome Project-style resting-state functional MRI at 7 Tesla using radiofrequency parallel transmission. NeuroImage 184, 396–408. https://doi.org/10.1016/j.neuroimage.2018.09.038

185. Yacoub, E., Hu, X., 2001. Detection of the early decrease in fMRI signal in the motor area. Magn Reson Med 45, 184–190. https://doi.org/10.1002/1522-2594(200102)45:2<184::aid-mrm1024>3.0.co;2-c

186. Yan, W.X., Mullinger, K.J., Brookes, M.J., Bowtell, R., 2009. Understanding gradient artefacts in simultaneous EEG/fMRI. NeuroImage 46, 459–471. https://doi.org/10.1016/j.neuroimage.2009.01.029

187. Yeh, M.-Y., Wu, C.W., Kuan, W.-C., Wei, P.-S., Wan, Y.-L., Wai, Y.-Y., Weng, H.-H., Liu, H.-L., 2013. Variations in BOLD response latency estimated from event-related fMRI at 3T: Comparisons between gradient-echo and Spin-echo. International Journal of Imaging Systems and Technology 23, 215–221. https://doi.org/10.1002/ima.22054

188. Yu, X., Qian, C., Chen, D., Dodd, S.J., Koretsky, A.P., 2014. Deciphering laminar-specific neural inputs with line-scanning fMRI. Nat Methods 11, 55–58. https://doi.org/10.1038/nmeth.2730

189. Zahneisen, B., Hugger, T., Lee, K.J., LeVan, P., Reisert, M., Lee, H.-L., Assländer, J., Zaitsev, M., Hennig, J., 2012. Single shot concentric shells trajectories for ultra fast fMRI. Magnetic Resonance in Medicine 68, 484– 494. https://doi.org/10.1002/mrm.23256

190. Zhang, W., Li, S., Wang, X., Gong, Y., Yao, L., Xiao, Y., Liu, J., Keedy, S.K., Gong, Q., Sweeney, J.A., Lui, S., 2018. Abnormal dynamic functional connectivity between speech and auditory areas in schizophrenia patients with auditory hallucinations. NeuroImage: Clinical 19, 918–924. https://doi.org/10.1016/j.nicl.2018.06.018

